# The solvent-excluded surfaces of water-soluble proteins

**DOI:** 10.1101/294082

**Authors:** Lincong Wang

## Abstract

The solvent-excluded surface (SES) of a protein is determined by and in turn affects protein-solvent interaction and consequently plays important roles in its solvation, folding and function. However, accurate quantitative relationships between them remain largely unknown at present. To evaluate SES’s contribution to protein-solvent interaction we have applied our accurate and robust SES computation algorithm to various sets of proteins and ligand-protein interfaces. Our results show that each of the analyzed water-soluble proteins has a negative net charge on its SES. In addition we have identified a list of SES-defined physical and geometrical properties that likely pertain to protein solvation and folding based on their characteristic changes with protein size, their differences between folded and extended conformations, and their correlations with known hydrophobicity scales and with experimentally-determined protein solubility. The relevance of the list of SES-defined properties to protein structure and function is supported by their differences between water-soluble proteins and transmembrane proteins and between solvent-accessible regions and ligand-binding interfaces. Taken together our analyses reveal the importance of SES for protein solvation, folding and function. In particular the universal enrichment of negative charge and the larger than average SES area for a polar atom on the surface of a water-soluble protein suggest that from a protein-solvent interaction perspective to fold into a native state is to optimize the electrostatic and hydrogen-bonding interactions between solvent molecules and the surface polar atoms of a protein rather than to only minimize its apolar surface area.

## 1 Introduction

Protein-solvent interaction is believed to contribute largely to the solvation, folding and structure of a water-soluble protein [1, 2, 3, 4, 5] and plays an important role in its function such as ligand binding [6]. However it is challenging to quantify such contributions [7, 8] using either experimental approach [9, 10] or theoretical model [11, 12, 13, 14, 15] or molecular dynamic (MD ^1^) simulation [16, 17] or structural information [18, 19]. For example, due to the difficulty to evaluate protein-solvent interaction it is not clear at present how evolution has optimized the surfaces of naturally-occurring water-soluble proteins to make them best adapted to aqueous solvent. Clues to possible adaptation may be found through a systematic and detailed analysis of the surfaces of different types of proteins with known structures. There exist three mathematical models for protein surface called respectively van der Waals (VDW) surface, solvent-accessible surface (SAS) [20, 21] and solvent-excluded surface (SES) [22, 23]. A SES is a two-dimensional (2D) manifold impenetrable to solvent molecules. In other words a SES defines a 2D boundary that seals off the interior of a protein from direct contact with solvent molecules [24]. The SES of any molecule consists of three different types of 2D patches: convex spherical polygons on a set of solvent-accessible atoms, saddle-shaped toroidal patches each of them defined by a pair of accessible atoms^2^ and concave spherical patches each of them determined by a triple of accessible atoms. In the past predominately SAS and to a much less degree SES have been extensively investigated mainly at residue-level for their roles in protein solvation, folding, stability and function [25, 26, 27, 21, 28, 29, 30, 31, 32, 9, 14]. For example it has been well documented that polar (hydrophilic) residues especially the charged ones prefer to be on the surface of a water-soluble protein while apolar (hydrophobic) residues are generally buried inside [33]. Further efforts have been made to establish quantitative relationships between SAS area and solvation free energy. For example, the free energies (∆*G*_solv_s) of the transfer of either organic compounds or small peptides between aqueous solvent and nonpolar solvents have been fitted to a linear equation ∆G_solv_ = *Σ*_*i*_ *σ*_*i*_ *A*_*i*_ where *A*_*i*_ is the SAS area of atom *i* of either a compound or a peptide. The fitted *σ*_*i*_s are called atomic solvation parameters [28]. Though such an empirical equation has found wide applications in various implicit solvent models for representing the contributions of solvent to protein folding, structure and ligand binding [14, 34], the physics behind the fitted *σ*_*i*_s is not well understood. Furthermore, to the best of our knowledge no efforts have been made in the past to establish a quantitative relationship between SES and protein-solvent interaction through a comprehensive analysis of the SESs for different types of proteins and ligand-protein interfaces at atomic level and on a large-scale.

To examine SES’s contribution to protein-solvent interaction at atomic level, to identify plausible physics behind atomic solvation parameter and to obtain clues to SES’s optimization via evolution we have applied our accurate and robust SES computation algorithm to a set 𝕊 of 16,483 water-soluble proteins with high quality crystal structures, a set 𝕄_e_ of 1,314 structural models of extended conformations and a set of proteins whose solubilities have been determined experimentally. The SESs of 𝕊 and 𝕄_e_ are further compared with the SESs of the lipid-exposing regions of transmembrane proteins and the SESs of ligand-protein interaction interfaces where ligand is either lipid or DNA or protein. Our analysis is inspired by the observations that water as a protic solvent prefers anions over cations as its solutes and both the intermolecular^3^ hydrogen bonding and the VDW attraction between the surface atoms of a solute and solvent molecules contribute to protein-solvent interaction. The analyses especially the comparisons of the atomic SES areas and atomic properties among different types of proteins and between the surfaces of water-soluble proteins and ligand-protein interfaces have identified a list of SES-defined physical and geometrical properties that are likely to be important for protein solvation, folding and function. This paper focuses on SES’s contribution to protein solvation and folding through the analyses of a list of SES-defined properties over 𝕊 and 𝕄_e_ while our sequels will demonstrate SES’s importance to protein structure and function using as examples the characteristic SES-defined properties for protein-protein [35], lipid-protein and DNA-protein interaction interfaces.

Our analyses show that every structure in S has a negative net surface charge. For example, the charges per atom for all the *accessible* atoms in 𝕊 have an average of −2.90 × 10^−2^*e* (elementary charge) while the charges per atom for all the *buried* atoms in S have an average of +2.70 × 10^−2^*e*. This large difference in charge per atom confirms quantitatively and at atomic level the residue-level observation that polar residues especially the charged ones prefer to be on the surface of a water-soluble protein [33]. Interestingly we find that compared with charge only or area only SES-area weighted surface charge and charge density seem to be more relevant to protein-solvent interaction. This finding provides a plausible explanation to atomic solvation parameters.

Our analyses have identified several SES-defined geometrical properties pertinent to intermolecular hydrogen bonding interaction. Specifically we find that SES area per accessible *polar* atom is, on average, almost 2-fold larger than SES area per accessible *apolar* atom. In our definition (section S1 of the Supplementary Materials) a polar atom is capable of forming a hydrogen bond with other atoms while an apolar one may not. In addition though the total SES area *A*_*i*_ of all the accessible polar atoms of a water-soluble protein is, on average, 1.2-fold smaller than the total SES area *A*_*o*_ of its accessible apolar atoms, *A*_*i*_ decreases but *A*_o_ increases upon unfolding^4^. Thus *A*_o_ and *A*_*i*_ as well as the ratio of SES area per apolar atom over SES area per polar atom likely pertain to protein-solvent interaction. These findings confirm quantitatively and at atomic level the preference of polar residues on the surface of a water-soluble protein. They also support the importance of intermolecular hydrogen bonding to protein solvent interaction [36] and may provide an alternative explanation [37] to some phenomena usually being associated with hydrophobic effect.

It is widely accepted that hydrophobic effect is the driving force for protein folding [2, 3, 7, 38, 39]. However, the quantitative contributions of hydrophobic effect to protein folding and PPI remain controversial [37]. One reason is that it has been difficult to evaluate the hydrophobic interaction between a folded water-soluble protein and solvent molecules since the protein surface is amphipathic. For an apolar solute it has been assumed that the intermolecular VDW attraction between the solute and aqueous solvent molecules is important for its solvation [39, 40]. Along this line of thinking we have identified a SES-defined geometrical property called concave-convex ratio *r*_*cc*_ that likely pertains to protein-solvent interaction. Our analysis shows that for a water-soluble protein the *r*_*cc*_ of an accessible *apolar* atom is, on average, 1.5-fold larger than the *r*_*cc*_ of a *polar* one. Most interestingly at residue-level *r*_*cc*_ correlates well with known hydrophobicity scales [41, 42, 43, 44]. These findings support the importance of intermolecular VDW attraction to the solvation of apolar atoms if we assume that the larger atomic *r*_*cc*_ is the stronger the VDW attraction between a protein surface atom and solvent molecules. These findings could also mean that the larger *r*_*cc*_ is, the less disruption to water’s hydrogen-bonded network [7].

The relevance to protein-solvent interaction and protein function of the list of SES-defined physical and geometrical properties is further supported by (a) their well-defined changes with protein size, (b) the differences between their values for folded proteins and for extended conformations, (c) the differences between their values for water-soluble proteins and for ligand-protein interfaces, and (d) the correlations between these properties and experimentally-determined solubility. From our large-scale analysis we hypothesize that the optimization of protein-solvent interaction through natural selection has been achieved via (1) the universal enrichment of negative surface charge, (2) the increased surface area for a surface polar atom for optimal hydrogen bonding with water molecules with minimal disruption to water’s hydrogen-bonded network, and (3) the increased concave-convex ratio for a surface apolar atoms for either stronger VDW attraction with water molecules or less disruption to water’s hydrogen-bonded network or both. This hypothesis is consistent with the observation that some of these SES-defined properties for de novo designed water-soluble proteins differ largely from those for naturally-occurring ones. It seems to us that a paradigm shift may be needed in the study of protein folding by taking a more balanced view of surface charge and side chain hydrophobicity since from a solvation perspective to fold into a native state is to optimize both the surface charges and the SES areas of the accessible *polar* atoms of a water-soluble protein rather than to only minimize the total SES area of its exposed *apolar* atoms.

## 2 Materials and Methods

In this section we first describe the data sets used in the analysis and then briefly present SES computation. Finally we define a list of SES-defined physical and geometric properties that likely pertain to protein-solvent interaction.

### 2.1 The data sets

We have downloaded from the PDB a non-redundant set of 25, 729 crystal structures of water-soluble proteins each has at most 70% sequence identity with any others, a resolution ≤ 3.5Å and a *R*-factor ≤ 27.5%. In this set each monomeric protein has > 800 atoms (with protons) and each multimer > 1, 000 atoms. This set excludes hyper-thermophilic, anti-freeze, membrane and nucleic acid binding proteins in order to minimize other structural features that may affect protein-solvent interaction. A prepossessing step that requires that no structures have > 5% missing atoms and no structures include bound compounds with > 20 heavy atoms reduces the number of structures to 16, 483. This set of structures is denoted as 𝕊 and is used as the representatives of water-soluble proteins. Set 𝕊 has the number of atoms ranging from 833 to 171, 552 and includes a set 𝕄 of 8, 974 monomeric proteins with 833 to 44, 200 atoms. Out of 𝕄 we select a subset 𝕄_f_ of 1,314 structures (section S2 of the Supplementary Materials) with 1, 004 to 10, 297 atoms that have coordinates for every residue, no bound compounds with > 5 atoms and < 0.2% missing atoms. Set 𝕄_f_ is used to represent water-soluble proteins in native state for the quantification of the changes in SES-defined properties upon unfolding. The corresponding model structures in unfolded state are a set of extended and energy-minimized conformations 𝕄_e_ generated by CNS [45] using the amino acid sequences in 𝕄_f_.

### 2.2 The preprocessing of PDB files for SES computation

The PDB files are preprocessed as follows for SES computation. Protons are first added using the program REDUCE [46] to any PDB structure that lacks their coordinates and the protonated structures are then processed by our structural analysis and visualization program. A graph with atom as node and bond as edge is first constructed for each of the 20 naturally-occurring amino acid residues, HSD, HSP and protonated ASP and GLU residues using Charmm atom nomenclature [34]. A molecule graph is then built for a whole protein by adding an edge for each peptide bond. For atoms with more than one conformation, only their first forms are selected for SES computation. Next any gap (a residue with no experimental coordinates) in a protein chain is identified and the percentage of missing atoms in each structure is computed by a comparison of the number of the nodes in the protein molecule graph with the number of atoms that have coordinates in the PDB file. Charmm force field parameters (e.g. Charmm partial charges) [34] are assigned to individual or a subset of atoms using a protein molecule graph. Only protein atoms are included in SES computation.

### 2.3 SES computation

A SES is composed of three types of areas: a spherical polygon area *a*_*s*_(*i*) on the surface of a solvent-accessible atom *i*, a patch area *a*_*t*_(*i, j*) on a toroid defined by two atoms *i, j* and a spherical polygon area *a*_*p*_(*i, j, k*) on the surface of a probe whose position is determined by three atoms *i, j, k.* The SESs and areas by our algorithm have higher accuracy than those by MSMS [47] due in part to the analytic computation of all the intersecting arcs among the probes, the accurate treatments of various cases of probe-probe intersections and no modifications to atomic radii [24]. In this study we set the probe radius to 1.4Å except for set 𝕄 over which SESs are computed twice using respectively 1.4Å and 1.2Å. The SESs with 1.2Å radius are compared with those with 1.4Å to see how probe radius affects area and SES-defined physical and geometrical properties^5^.

### 2.4 SES-defined physical and geometrical properties

A list of physical and geometrical properties have been defined using atomic SES to evaluate their possible contributions to protein solvation and function. These SES-defined properties are inspired by the observations that water as a protic solvent prefers anions over cations, and that both the hydrogen bondings between solvent molecules and the polar atoms of a solute and the VDW attractions between solvent molecules and its apolar atoms contribute to its solvation. Their definitions rely on atomic SES area. However except for atomic concave-convex ratio each of the other properties is defined over a specific set of atoms.

To each accessible atom *i* we assign an atomic SES area *a(i):*

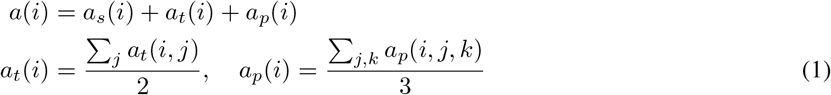

where *a*_*s*_(*i*), *a*_*t*_(*i*) and *a*_*p*_(*i*) are respectively the accessible, toroidal and probe areas for atom *i.* From *a*_*s*_(*i*) and *a*_*p*_(*i*) we define a concave-convex ratio *r*_*cc*_(*i*) for atom *i* to estimate its local ruggedness and 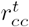 for a set of accessible atoms T to represent the average ruggedness of the surface formed by them. For example the *r*_*cc*_ for the set of accessible atoms belonging to a single residue is called residue *r*_*cc*_.

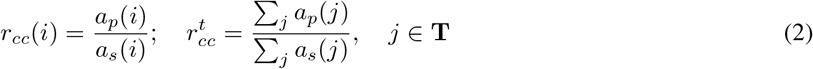

On the set **A** of accessible atoms of a protein we define as follows its SES area *A,* net surface charge *Q*_A_, surface charge density Σ_A_, average-partial charge (charge per atom) *ρ*_A_, average-atomic area (area per atom) *η*, and surface atom density (number of atoms per area) *ν.*

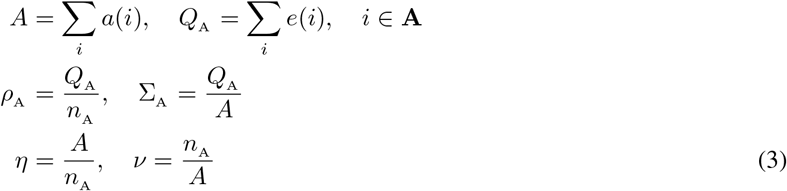

where *n*_A_ = |**A**| is the number of accessible atoms and *e*(*i*) the Charmm partial charge for atom *i* [34]. By Eq. (3) we have 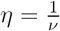. On the set of buried atoms **B** in a protein we define its net charge 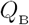 and average-partial charge 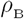.

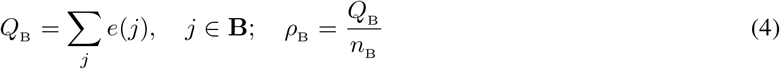

where *n*_B_ = |**B**| is the number of atoms in **B**. The net charge *Q,* and average-partial charge *ρ* for a *whole* protein are defined as follows.

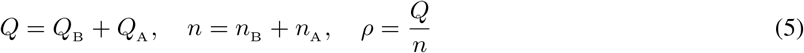

where *n* = |**N**| is the total number of atoms in a protein and set **N** = **A** ∪ **B** includes all of its atoms. Area-weighted surface charge *q*_*s*_ and area-weighted surface charge density *σ*_*s*_ are defined as follows to represent simultaneous contributions of surface charge and area to protein-solvent interaction.

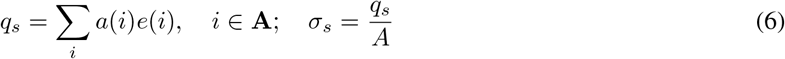

To distinguish the different contributions to protein-solvent interaction between accessible *polar* atoms and accessible *apolar* atoms we divide **A** into two different subsets, set **A**_o_ of apolar atoms and set **A**_i_ of polar atoms, that is, **A** = **A**_o_ ∪ **A**_i_. The accessible atoms in A_i_ are either hydrogen bond donors or acceptors as specified in Charmm force field [34] while those in **A**_o_ include the rest. On both **A**_i_ and **A**_o_ we define as follows their respective SES areas *A*_o_, A_*i*_ and their ratio *A*^*oi*^, average-atomic areas *η*_*i*_ and *η*_*o*_ and their ratio *R*^*io*^, concave-convex ratios 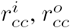 and their ratio 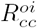

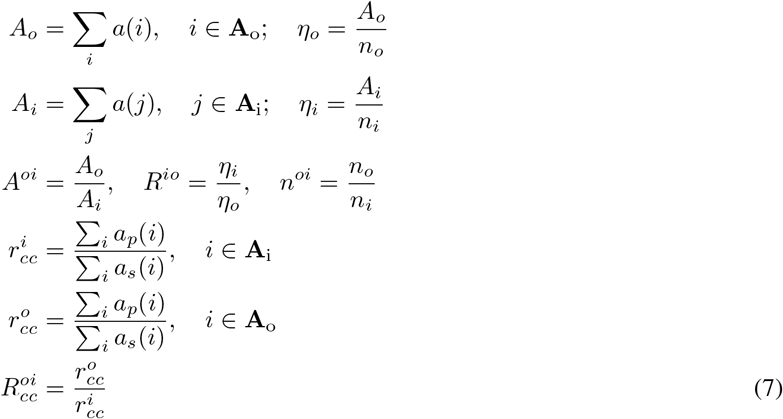

where *n*_*o*_ = |**A**_o_| and *n*_*i*_ = |**A**_i_| are respectively the numbers of atoms in **A**_o_ and **A**_i_, and *n*^*oi*^ is their ratio. The SES areas *A*_*i*_ and *A*_*o*_ are called respectively *the polar surface area* and *the apolar surface area* of a protein.

## 3 Results and Discussion

In this section we first briefly describe the processing of PDB structure files. We then present the analyses of the list of SES-defined properties on set 𝕊, 𝕄_f_ and 𝕄_e_, and discuss their relevance to protein solvation and folding. The importance of this list of properties to protein function is discussed in terms of their differences between 𝕊 and ligand-protein interaction interfaces where ligand is either lipid or DNA or protein. Overall in terms of SES-defined properties the differences between 𝕊 and 𝕄_f_ are statistically insignificant while the differences between 𝕄_f_ and 𝕄_e_ are relatively large and the differences between ligand-protein interfaces and 𝕊 are substantial.

### 3.1 The processing of PDB structure files

In order to eliminate as much as we could other factors that may interfere with our SES analysis, we have applied a list of strict criteria to ensure that the sets of analyzed structures have good structural qualities and whose surfaces are representatives of water-soluble proteins. Both the SES and the structure of any protein that has a SES-defined property in the upper or lower 1.0% of its distribution over 𝕊 are inspected visually using our structural analysis and molecular visualization program to make sure that the PDB file has been properly processed. Any PDB file that could not be correctly processed by our program is removed from further analysis. Such an outlier is further checked against literature to ensure it is not one of hyperthermophilic, anti-freeze, membrane and DNA-binding proteins.

### 3.2 The surface charges of water-soluble proteins

Previous studies on protein surfaces mainly SAS and VDW surfaces and to a much less extent SESs have shown that polar residues especially charged ones prefer to be on the surface of a water-soluble protein [33]. In principle protein-solvent interaction is electrostatic in nature^6^ [16, 48]. In theory surface charge and dipole moment are closely related to protein solvation [11, 12, 13, 14, 15]. Inspired by the importance of electrostatic interaction for solvation especially by the observation that water as a protic solvent prefers anions over cations we first analyze the differences in charge between accessible atoms and buried atoms. As shown in Fig. 1 we discover that each of the 16,483 proteins in 𝕊 has a negative net charge (negative *Q*_A_ and *ρ*_A_) for its accessible atoms and a positive net charge (positive *Q*_B_ and *ρ*_B_) for its buried atoms. Most strikingly the difference between the average *ρ* for all the sets of the accessible atoms in 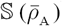^7^ and the average *ρ* for all the sets of the buried atoms in 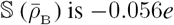, and the ratio 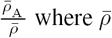 is the average net charge for all the atoms in a protein is 19.33, equivalent to a 19-fold difference in negativity between the accessible atoms and all the atoms. In addition *Q*_A_ increases with protein size^8^ via a well-fitted power law and the enrichment in negativity is apparent for the folded (native) structures in 𝕄_f_ when compared with the extended conformations in 𝕄_e_ (Table 1). In stark contrast with the average 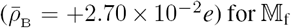, the average 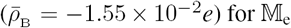 is negative while the average 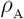 for 𝕄_e_ is more than 100-fold less negative than that for 𝕄_f_ (Table 1). The negativity of 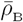 for 𝕄_e_ is due mainly to the buried backbone nitrogen and oxygen atoms. Furthermore, the 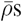 for the buried atoms in PPI interfaces [35] and DNA-protein interfaces are both *positive,* and the 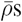 becomes less negative for the lipid-exposing regions of transmembrane proteins and for the surface atoms that become buried upon ligand bindings.

**Figure 1:**
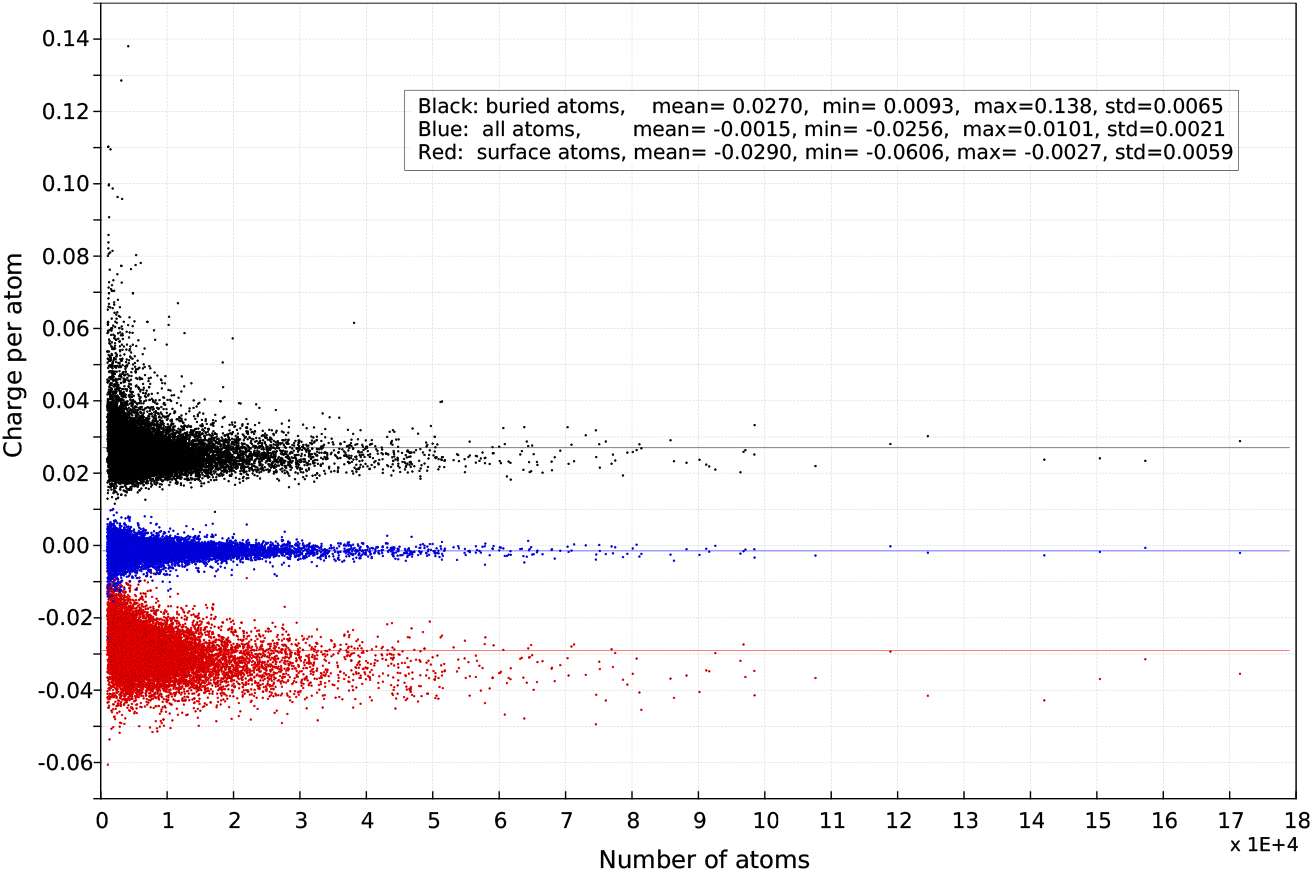
The average-partial charges *ρ*_A_, *ρ*_B_ and *ρ* for 𝕊. They are colored respectively in red, black and blue and their means (*µs*) are respectively −0.0290*e*, 0.0270*e* and −0.0015*e* per atom. The ratio between the *µ*s for *ρ*_A_ and *ρ* is 19.333. The x-axis is the number of atoms in a structure. The y-axis is the average-partial charge of a structure with a unit of *e* per atom. All the plots in this paper are prepared using an in-house 2D plot program written in Qt/C++.

Another SES-defined electrical property is surface charge density Σ. As shown in Fig. 2 the three surface charge densities, 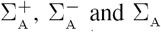, for the extended conformations in 𝕄_e_ differ largely from those for 𝕄_f_. For the native structures in 𝕄_f_, 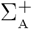 increases while both 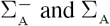 decrease with protein size. If we fit the Σ_A_ s for 𝕄_f_ to a power law, Σ = *an*^*b*^ + c, where *n* is number of atoms (protein size), then the fitted parameter *c* = −5.00 × 10^−3^ is much more negative than the Σ_A_ average 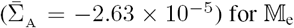. The extended conformations in 𝕄_e_ likely deviate from the real unfolded states existent in a typical experimental setting [49] and thus their SES-defined properties differ from those for a genuine unfolded state. However the large differences in Σs between 𝕄_f_ and 𝕄_e_ support at least qualitatively the relevance of net surface charge density to solvation and folding. In addition as shown in Figs. 11(d) and 12(d) there exist good correlation between Σ_A_ and experimentally-determined solubility. As to be expected, more negative Σ_A_ value a protein has better solubility in aqueous solution.

**Table 1:** The average-partial charges, *ρ*_A_, *ρ*_B_ and *ρ,* of folded structures and extended conformations. The two values in each cell are respectively mean (average) and standard deviation with a unit of *e.* The differences in 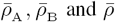 between the SESs for set 𝕊 and its subset 𝕄_f_ are rather small.

| | $\rho_A$ | $\rho_B$ | $\rho$ |
| --- | --- | --- | --- |
| $\mathbb{M}_f$ | $-2.73 \times 10^{-2} / 0.55 \times 10^{-2}$ | $+2.7 \times 10^{-2} / 0.64 \times 10^{-2}$ | $-1.2 \times 10^{-3} / 2.0 \times 10^{-3}$ |
| $\mathbb{M}_e$ | $-1.65 \times 10^{-4} / 20.0 \times 10^{-4}$ | $-1.55 \times 10^{-2} / 3.0 \times 10^{-2}$ | $+3.68 \times 10^{-4} / 0.24 \times 10^{-4}$ |

**Figure 2:**
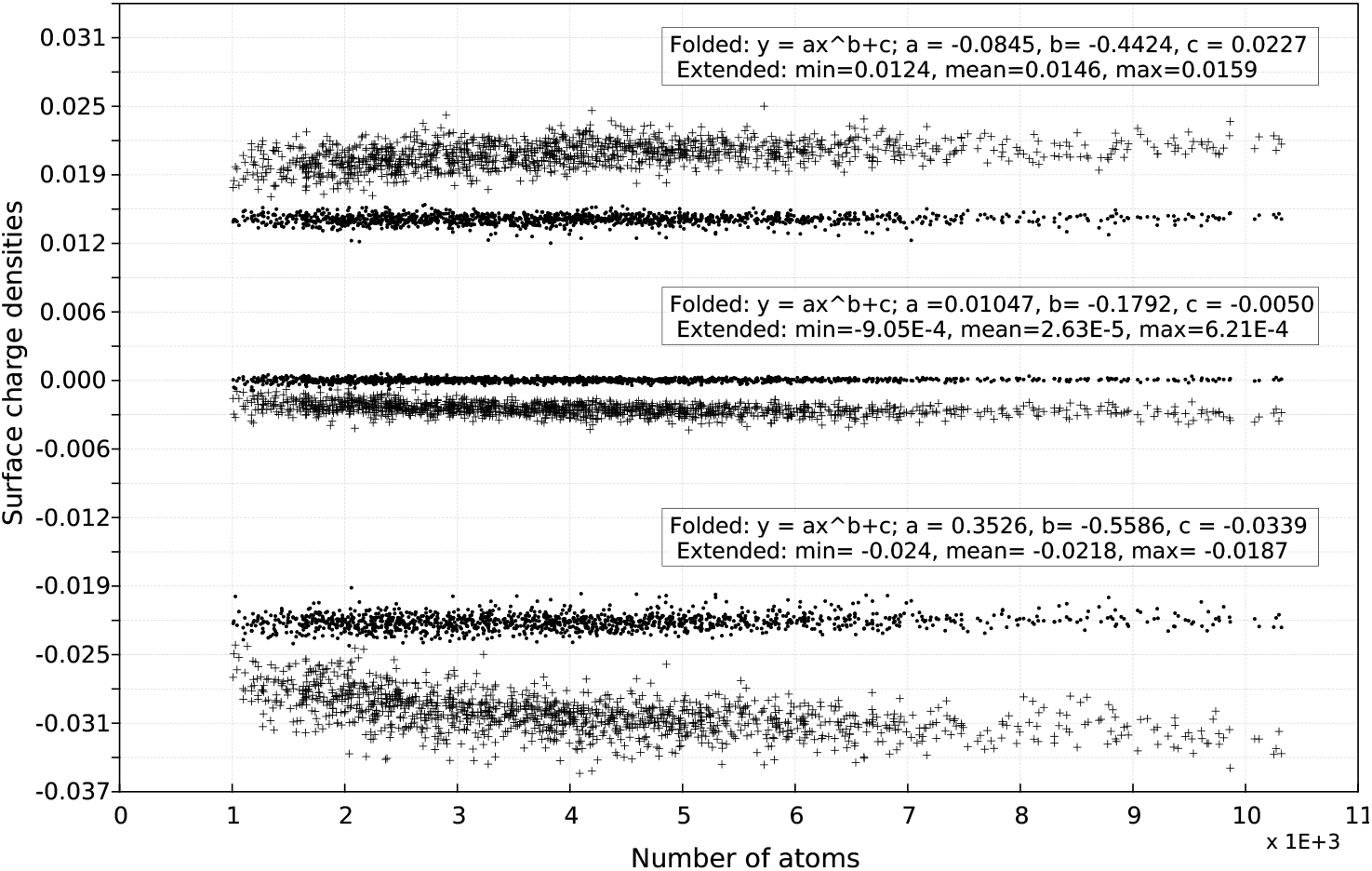
The surface charge densities of folded structures in 𝕄_**f**_ vs extended conformations in 𝕄_**e**_. The three surface charge densities 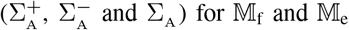 are shown respectively as filled circles and crosses. To compute 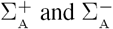 set **A** is first divided into two subsets **A**^+^ and **A**^−^ where **A**^+^ is composed of all the accessible atoms with a positive partial charge while all the accessible atoms with a negative partial charge belong to **A**^+^. Then 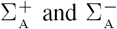 are computed as 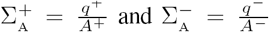 where 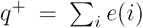 and 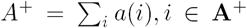 and 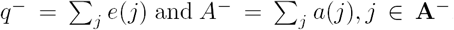. The inserts list the minimums, maximums and means for the three densities over 𝕄_e_ and the three fitted power laws for 𝕄_f_. The x-axis is the number of atoms in a structure. The y-axis is surface charge density in *e* / Å^2^.

However, as shown in Figs. 1, 2 and Fig. 5 of section 3.3, neither *ρ* nor Σ nor *η* (area per atom) changes linearly with protein size (*n*) and the distributions around their means are not symmetrical especially for small-sized proteins. The non-uniformity implies that none of them alone could provide a proper description to protein-solvent interaction because its strength is expected to be statistically independent of *n.* In contrast to *ρ,* Σ and *η,* area-weighted surface charge (*q*_*s*_) changes almost linearly with *n* and area-weighted surface density (*σ*_*s*_) is almost independent of *n* (Fig. 3). In addition the distribution around the mean for *σ*_*s*_ is rather symmetrical as indicated by a very small difference between its mean and median even for small-sized proteins. More interestingly each of the 16,483 proteins in 𝕊 has a negative *σ*_*s*_ (Fig. 3). In addition as shown in Table 2 the ratio between the *σ*_*s*_ for a folded structure in 𝕄_f_ and the *σ*_*s*_ for a corresponding extended conformation in 𝕄_e_ has an average of 1.57. Furthermore, the *σ*_*s*_s for the lipid-exposing atoms of transmembrane proteins and for the interface atoms that become buried upon ligand-binding all become less negative. Thus the three SES-defined area-weighted properties, 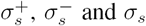, will likely provide a more balanced description to protein solvation, folding and function. In particular the expression for area-weighted surface charge 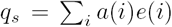 resembles the expression for atomic solvation parameters. Thus atomic solvation parameter *σ*_*i*_ is possibly related to partial charge *e(i).*

**Table 2:** The ratios of the 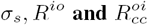 values of a folded structure over those of an extended conformation. The four values in a cell are respectively the minimum, mean, maximum and standard deviation of the distribution over 𝕄_e_ and 𝕄_f_ of a ratio.

| $\sigma_s$ ratio | $R^{io}$ ratio | $R_{cc}^{oi}$ ratio |
| --- | --- | --- |
| 1.17, 1.57, 2.01; 0.123 | 1.0, 1.22, 1.42; 0.062 | 0.91, 1.31, 1.69; 0.137 |

In summary our large-scale analysis shows that folding into a native state in aqueous solution turns a water-soluble protein into a capacitor with a positive net charge buried inside and a negative net charge on its SES (the outer surface of the capacitor) to maximize its electrostatic attraction to the solvent [50]. In other words, a water-soluble protein behaves, on average and as far as surface charge is concerned, as a micelle with an exterior formed predominately by atoms with negative partial charges and an interior composed of mainly atoms with positive partial charges. By extension there must exist a 2D manifold (the inner surface of the capacitor) inside a water-soluble protein that encloses a set of atoms with zero net charge. A model of alternative layers of negative and positive charges has been alluded before in MD simulation [51].

**Figure 3:**
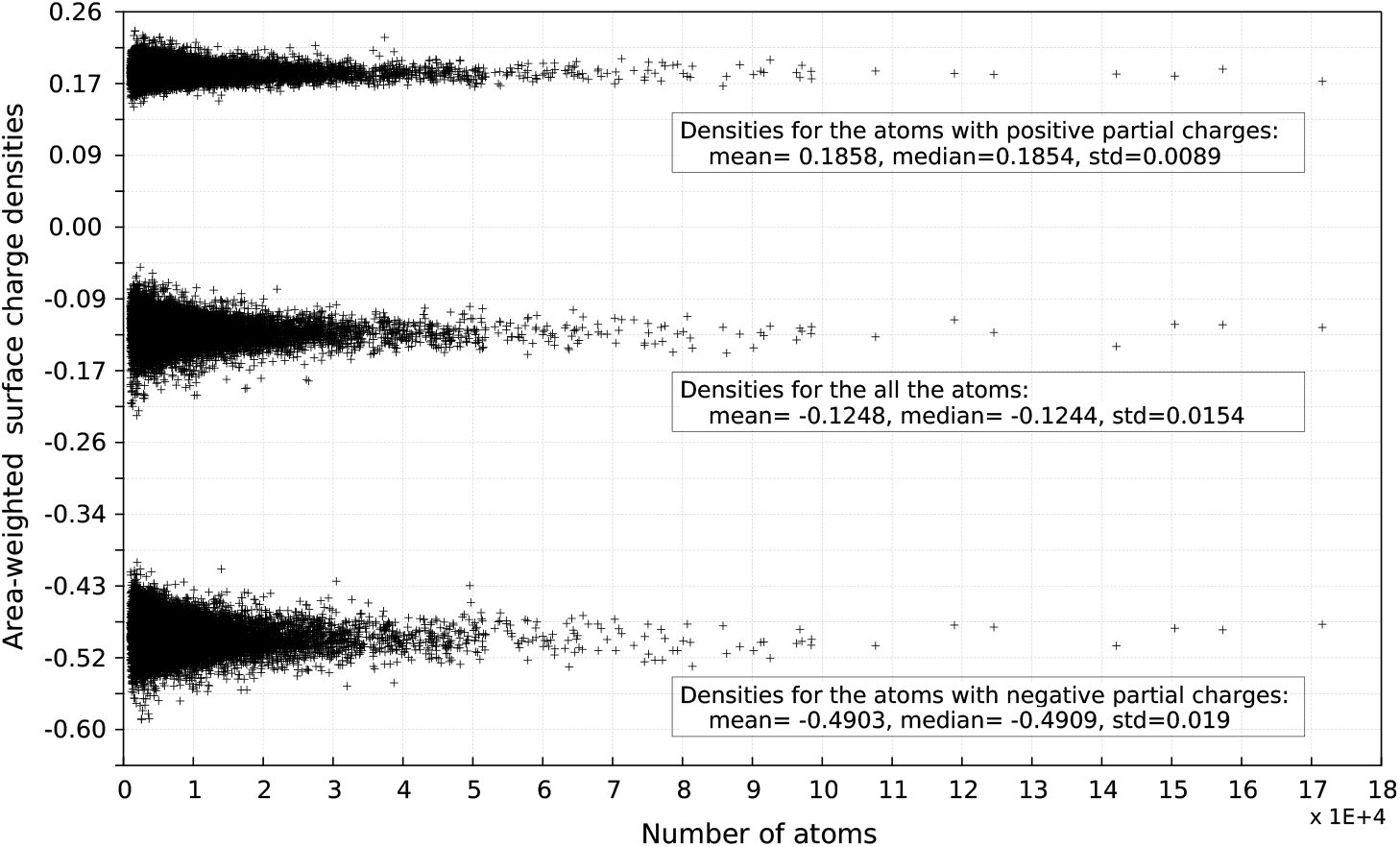
Area-weighted surface charge densities 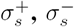 and *σ*_*s*_ for 𝕊. The 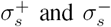 are defined as 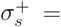 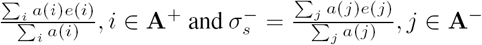. The three inserts list their respective means, medians and standard deviations. The x-axis is the number of atoms in a structure. The y-axis is area-weighted surface charge density in *e.*

### 3.3 Accessible polar and apolar atoms and their SES areas

Previous structural analyses [26, 41, 30, 33] have found that polar residues prefer to be on the surface of a water-soluble protein while apolar ones are likely to be buried inside. Such preferences are often cited as one piece of evidence for the importance of hydrophobic effect to the folding of a water-soluble protein. With the assignment of a SES area to an individual atom and the division of the set of accessible atoms into polar and apolar ones it is possible to quantify such preferences at atomic level using SES-defined physical and geometrical properties. The ratio of the number of accessible apolar atoms over that of polar atoms, *n^oi^,* is a property that could possibly quantify at atomic level the preference of polar atoms on the surface of a water-soluble protein. However 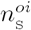 average for 𝕊 is 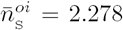, and *n*^*oi*^ increases very slowly with protein size *n* when *n <* 10, 000 and remains essentially the same when *n* > 10, 000 (Fig. S1 of the Supplementary Materials). It means that for the water-soluble proteins in 𝕊 the numbers of apolar atoms are on average more than 2-fold larger than the numbers of polar atoms. As with *n*^*oi*^ the SES-defined property *A*^*oi*^ has an average 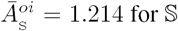 and on average the *A*^*oi*^s do not change with protein size (Fig. S2 of the Supplementary Materials). Thus the set of accessible apolar atoms in a typical water-soluble protein still has larger SES area than its set of accessible polar atoms. On the other hand the 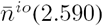 for the buried atoms in 𝕊 is 13.7% larger than the 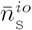 for 𝕊 (Fig. S1 of the Supplementary Materials). In addition the 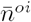 for 𝕄_e_ increases to 2.457 and the 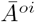 for 𝕄_e_ increases to 1.570. Furthermore both *A*_*i*_ and *A*_*o*_ decrease upon folding though *A*_*o*_ ≥ *A*_*i*_ remains to be true. Thus as been shown before at residue level [26, 41, 30, 33] folding into a native state indeed reduces both the number and the area of surface apolar atoms. A SES-defined property that could more directly quantify the previously-documented preferences for polar residues is area per atom *η.* As shown in Fig. 4 the ratio, 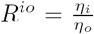, for 𝕊 ranges from 1.451 to 2.555 with 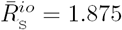. In other words, a polar atom has, on average, 1.875-fold larger SES area than an apolar atom. More interestingly only three structures (2ouw, 3qva and 4z0m) in 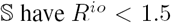. In addition the average *R*^*io*^ for 𝕄_e_ is 1.567, a 17.8% smaller than 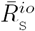. Furthermore though both *η*_*i*_ and *η*_*o*_ decrease upon folding the reduction in *η*_*i*_ is smaller than that in *η*_*o*_ (Fig. 5). One possible explanation for a large 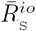 value is the importance to protein-solvent interaction of the intermolecular hydrogen bonding between accessible polar atoms and solvent molecules [36]. A large SES area for an accessible polar atom is likely to be favorable for optimal hydrogen bonding. The inter-atomic distance between two hydrogen-bonded atoms is smaller than the summation of their respective VDW radii. The larger SES area a polar atom has, the less likely a solvent molecule clashes with its neighboring protein atoms and less likely perturbs water’s hydrogen-bonded network when they form an optimal intermolecular hydrogen bond.

The relevance to protein solvation and function of the four SES-defined properties, *n*^oi^, *η*, *A*^*oi*^ and *R^io^,* is supported by the following observations. The 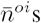 for the lipid-exposing regions of transmembrane proteins, PPI interfaces and lipid-protein interfaces are all larger than 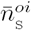 while the 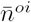 for DNA-protein interfaces is smaller than 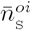. As with 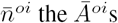 for lipid-exposing regions, PPI interfaces and lipid-protein interfaces are all larger than 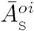. Significantly as shown in Figs. 11(b) and 12(b) *A*^*oi*^ correlates well with experimentally-determined protein solubility. However in contrast to 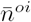 and 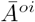, the 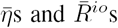 for the lipid-exposing regions of transmembrane proteins, lipid-protein interfaces, DNA-protein interfaces and PPI interfaces are all smaller than the respective 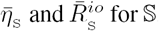. Furthermore, as shown in Fig. 4 and Table 3 the seven structures in 𝕊 with 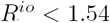 are either PSI targets with unknown functions or proteins that seem to interact with lipids in some fashions. Their *R*^*io*^ values are close to those for 𝕄_e_ and to those for PPI interfaces [35]. On the other hand, four (three ferredoxins and one flavodoxin) of the nine structures in 𝕊 that have their 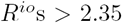 (Fig. 4 and Table 4) are involved in electron-transfer, two are DNA mimics, the other two are putative hemolysins, and 5cwh is a de novo designed protein [52]. The contrast between the SES of a protein with a large *R*^*io*^ and the SES of a protein with a small *R*^*io*^ is visually detectable: as shown in Fig. 6 the former has more largely-exposed *polar* atoms per SES area while the latter has more largely-exposed *apolar* atoms per SES area.

**Table 3:** The seven structures in S with *R*^*io*^ < 1.54. The nine SES-defined physical and geometrical properties are three atomic partial-charges *ρ*_A_, *ρ*_*B*_ and *ρ,* three SES area-weighted surface charge densities 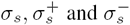, and three ratios 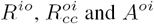. The second column is the total number of atoms in a structure. The last row lists their averages for 𝕊. There exists no correlation between the seven smallest *R*^*io*^s and the three atomic partial charges *ρ*_A_ *, ρ*_B_ and *ρ.*

| pdbid | $n$ | $\rho_A$ | $\rho_B$ | $\rho$ | $\sigma_s$ | $\sigma_s^+$ | $\sigma_s^-$ | $R^{io}$ | $R_{cc}^{oi}$ | $A^{oi}$ |
| --- | --- | --- | --- | --- | --- | --- | --- | --- | --- | --- |
| 2qsk | 1269 | -0.02375 | 0.04558 | 0.00269 | -0.09007 | 0.17609 | -0.44169 | 1.50466 | 0.95139 | 1.31158 |
| 2rcd | 7809 | -0.03372 | 0.03441 | -0.00126 | -0.11634 | 0.17715 | -0.46242 | 1.51725 | 1.01543 | 1.46297 |
| 2rfr | 2288 | -0.03078 | 0.04303 | -0.00084 | -0.11759 | 0.18211 | -0.47997 | 1.50019 | 1.04953 | 1.52315 |
| 2ouw | 3989 | -0.02920 | 0.03028 | -0.00057 | -0.11053 | 0.17059 | -0.44484 | 1.46541 | 0.97992 | 1.73937 |
| 3qva | 6109 | -0.03236 | 0.02721 | 0.00034 | -0.11655 | 0.19231 | -0.45482 | 1.48751 | 0.99860 | 1.35483 |
| 4qxl | 1704 | -0.01528 | 0.02510 | 0.00351 | -0.08379 | 0.21135 | -0.46661 | 1.52015 | 1.09481 | 1.13643 |
| 4z0m | 10090 | -0.03762 | 0.03145 | -0.00026 | -0.09640 | 0.18138 | -0.43458 | 1.45094 | 1.02202 | 1.77460 |
| $\mathbb{S}$ | | **-0.0290** | **0.0270** | **-0.0015** | **-0.125** | **0.186** | **-0.490** | **1.875** | **1.496** | **1.214** |

The enrichment of polar atoms, the enlargement of their total areas especially the large increase in SES area per polar atom on the SES of a water-soluble protein are consistent with the previous view that the hydrogen bonding interactions between surface polar atoms and solvent molecules contribute largely to protein solvation, folding and function. In addition there exist no or only weak correlations between SES-defined electrical properties such as *ρ*_*A*_ and σ_s_ and geometrical properties such as SES area, *A*^oi^ and *R*^*io*^ (section S6 of the Supplementary Materials). Furthermore the differences in SES area between polar surface atoms and apolar ones are in line with the heterogeneity of water motion in the first hydration shell. Thus from an evolutionary perspective it seems that the surfaces of naturally-occurring water-soluble proteins have evolved for best interaction with aqueous solvent through optimal intermolecular hydrogen bondings between surface polar atoms and solvent molecules. The importance of intermolecular hydrogen bondings to protein-solvent interaction may provide an explanation to hydrophobic effect [37].

**Figure 4:**
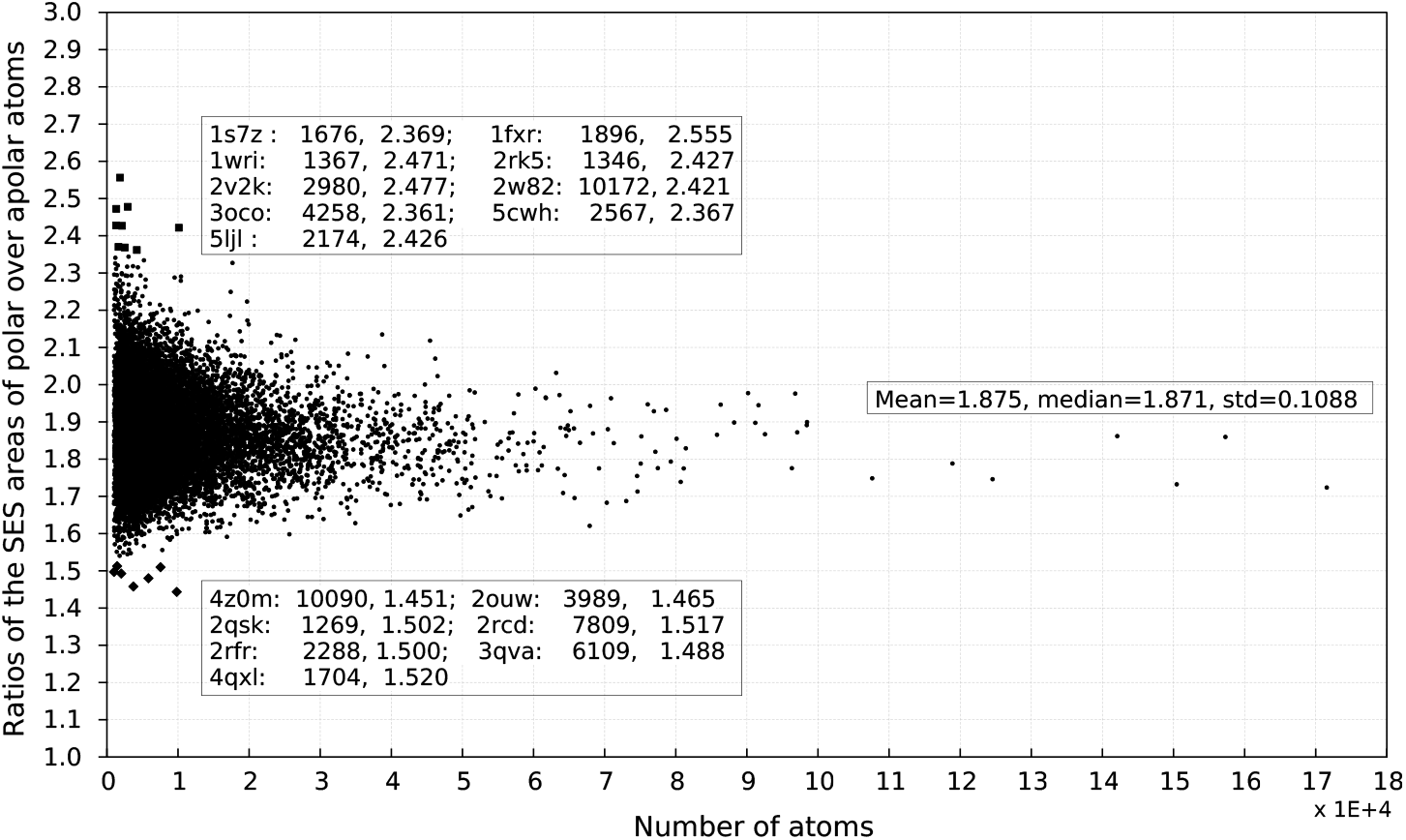
The ratio *R*^*io*^s for 𝕊. The middle insert lists *R*^*io*^’s mean, median and standard deviation for 𝕊. The top insert lists nine proteins that have *R*^*io*^ > 2.35 with their ratios depicted as filled squares. Among them are three ferredoxins (1fxr, 2v2k and 1wri) with the largest *R*^*io*^ values and a flavodoxin (5ljl), two DNA mimics (1s7z and 2w82), two putative hemolysins (2rk5 and 3oco) and one de novo designed protein (5cwh). The bottom insert lists seven proteins that have *R*^*io*^ < 1.54 with their ratios depicted as filled diamonds. Among them are three PSI targets (2rfr, 2ouw and 2rcd) with unknown functions, an antiviral lectin scytoririn (2qsk), a 5-hydroxyisourate hydrolase (3qva), a flagellar type III secretion operon (4qxl), and an isomerase (4z0m) that is involved in unsaturated lipid assimilation and has the smallest *R*^*io*^ value among all the proteins in 𝕊. The rest are depicted as filled circles. The x-axis is the number of atoms in a structure while the y-axis is *R*^*io*^.

**Table 4:** The nine structures in 𝕊 with *R*^*io*^ > 2.35. The nine SES-defined properties are the same as those in Table 3. The second column is the total number of atoms in a structure. The last row lists their averages for 𝕊. There exists only very weak correlation between the nine largest *R*^*io*^s and the three atomic partial-charges *ρ*_A_, *ρ*_B_ and *ρ.*

| pdbid | $n$ | $\rho_A$ | $\rho_B$ | $\rho$ | $\sigma_s$ | $\sigma_s^+$ | $\sigma_s^-$ | $R^{io}$ | $R_{cc}^{oi}$ | $A^{oi}$ |
| --- | --- | --- | --- | --- | --- | --- | --- | --- | --- | --- |
| 1fxr | 1896 | -0.05021 | 0.03033 | -0.01542 | -0.22582 | 0.17405 | -0.54620 | 2.55518 | 1.89535 | 1.00432 |
| 1s7z | 1676 | -0.04192 | 0.02347 | -0.01391 | -0.20103 | 0.18743 | -0.52662 | 2.36909 | 1.82902 | 1.09810 |
| 1wri | 1367 | -0.04121 | 0.02386 | -0.01070 | -0.20790 | 0.18142 | -0.54729 | 2.47127 | 1.82116 | 0.90102 |
| 2rk5 | 1346 | -0.03654 | 0.02625 | -0.00692 | -0.18591 | 0.19718 | -0.55358 | 2.42685 | 2.14083 | 0.86174 |
| 2v2k | 2980 | -0.04424 | 0.02832 | -0.01147 | -0.20535 | 0.16217 | -0.51384 | 2.47683 | 1.78641 | 1.06883 |
| 2w82 | 10172 | -0.04270 | 0.02490 | -0.01247 | -0.20160 | 0.18647 | -0.53317 | 2.42058 | 2.15347 | 0.99330 |
| 3oco | 4258 | -0.03349 | 0.02195 | -0.00625 | -0.17145 | 0.19731 | -0.52968 | 2.36075 | 1.99922 | 1.00332 |
| 5cwh | 2567 | -0.03282 | 0.03051 | -0.00674 | -0.15514 | 0.17671 | -0.54551 | 2.36745 | 2.37866 | 1.12571 |
| 5ljl | 2174 | -0.04801 | 0.02181 | -0.01249 | -0.22007 | 0.19293 | -0.55486 | 2.42575 | 1.92753 | 0.93006 |
| $\mathbb{S}$ | | **-0.0290** | **0.0270** | **-0.0015** | **-0.125** | **0.186** | **-0.490** | **1.875** | **1.496** | **1.214** |

**Figure 5:**
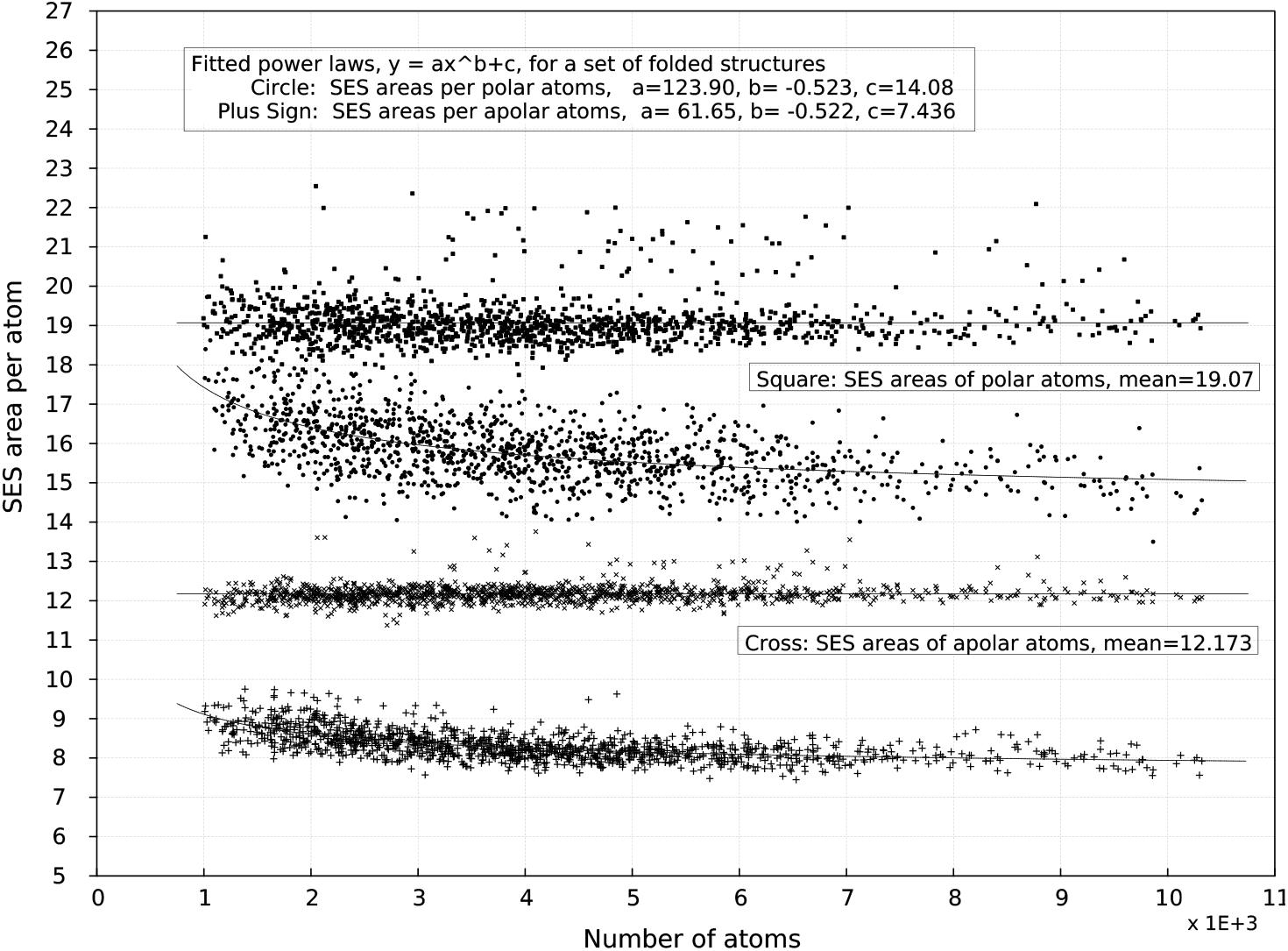
The *η*_*i*_s and *η*_o_s of the individual sets of accessible polar and apolar atoms in 𝕄_f_ and 𝕄_e_. The two curves represent respectively the fitted power laws for the *η*_*i*_s of the individual sets of accessible polar atoms (filled circles) in 𝕄_f_ and for the *η*_*o*_s of the individual sets of accessible apolar atoms (plus signs) in 𝕄_f_. The two lines indicate respectively the means for the *η*_*i*_s of the individual sets of accessible polar atoms (filled circles) in 𝕄_e_ and for the *η*_*o*_s of the individual sets of accessible apolar atoms (plus signs) in 𝕄_e_. Upon folding the reduction in *η*_*o*_ is estimated to be 38.9% = 100.0 × 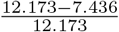 while reduction in *η*_*i*_ is only 26.2% = 100.0 × 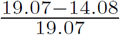. The x-axis is the number of atoms in a structure while the y-axis is either *η*_*i*_ or *η*_*o*_.

### 3.4 The SES geometry of polar and apolar atoms

One advantage of SES over SAS is that the former includes both convex and concave areas while the latter has only convex ones. With SES we could define a concave-convex ratio *r*_*cc*_ either for a single atom or over a set of accessible atoms such as the set of all the accessible atoms of a surface residue and the set of all the accessible atoms of a protein (Eqs. 2 and 7). To see the possible relevance of *r*_*cc*_ to protein-solvent interaction we have analyzed the *r*_*cc*_s for 𝕊, 𝕄_f_ and 𝕄_e_ as well as the *r*_*cc*_s for the lipid-exposing regions of transmembrane proteins and ligand-protein interaction interfaces. Both the 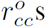 and the 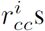 for 𝕊 increase with protein size via well-defined power laws. More relevantly their ratio 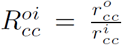 is independent of protein size and ranges from 0.951 to 2.833 with a mean of 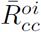 = 1.496 (Fig. 7). In fact except for four structures, 2qsk, 3vqj, 2ouw and 3qva, the 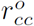 for each water-soluble protein in 𝕊 is larger than its 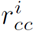. The relevance of *r*_*cc*_ and 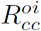 to protein solvation, folding and function is further supported by the following observations. Firstly, the 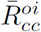 for 𝕄_e_ is 1.31-fold smaller than that for 𝕄_f_ (Table 2). Interestingly, compared with the *r*_*cc*_s for 𝕄_f_, the *r*_*cc*_s for 𝕄_e_ do not change with protein size and are several-fold smaller (Fig. 8). Secondly, the 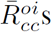 for the lipid-exposing regions of transmembrane proteins, lipid-protein interfaces, DNA-protein interfaces and PPI interfaces are all smaller than that for 𝕊. Particularly the 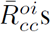 for the lipid-exposing regions of transmembrane proteins and lipid-protein interfaces are close to 1.0. Accordingly we expect that a protein that has a 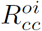 value close to 1.0 (Fig. 7 and Table 5) is likely either a peripheral membrane protein or a lipid-binding protein. For example, a previous experiment has shown that the expression in *E.coli* of an antiviral lectin scytoririn led to the accumulation of the expressed proteins in membrane [53]. Thirdly, the *r*_*cc*_s for PPI interfaces are several-fold smaller than those for 𝕊 [35]. And finally as shown in Table 7 and Fig. 9, the solvent-accessible residue *r*_*cc*_s correlate well with known hydrophobicity scales. There exists modest correlation between 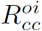 and *R*^*io*^ (section S6 and Fig. S6 of the Supplementary Materials) likely because both are defined in terms of **A**_i_ and **A**_0_ (Eqn. 7).

**Figure 6:**
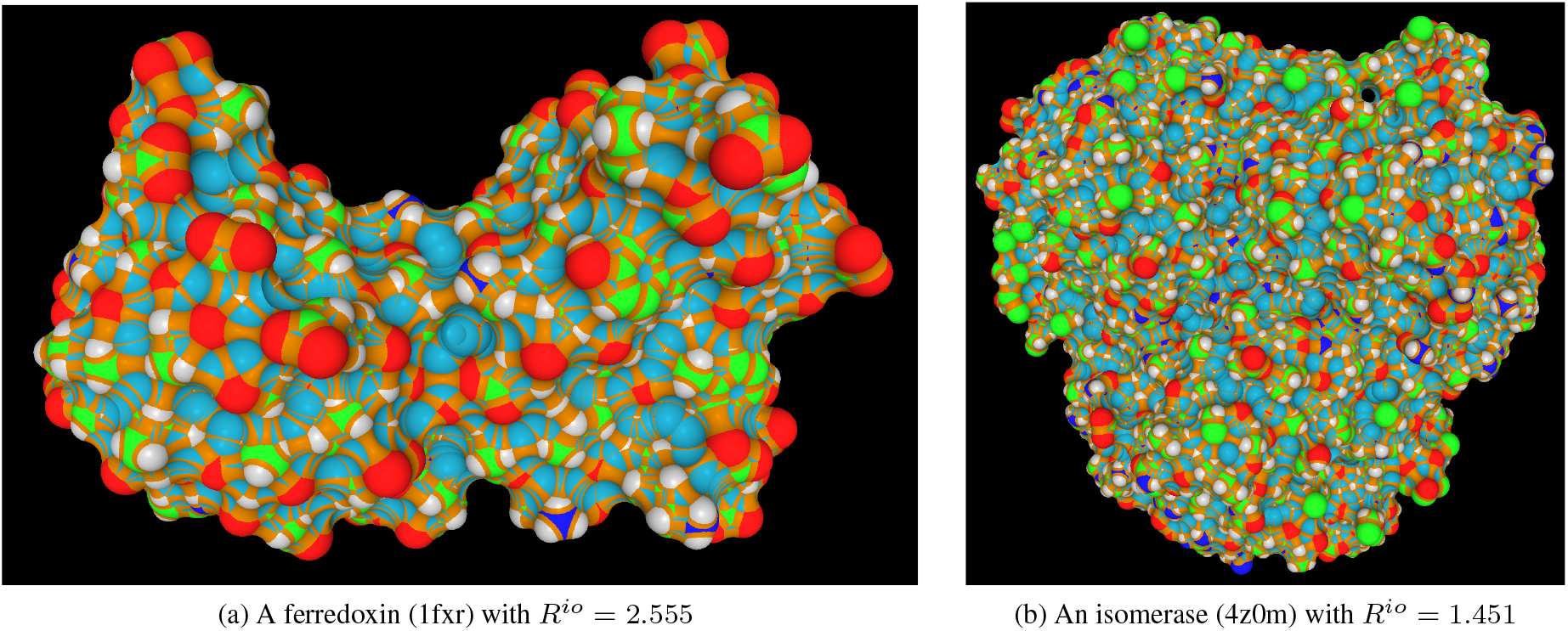
The SESs of the two proteins with the largest and the smallest *R*^*io*^ values among all the proteins in 𝕊. The accessible C, O, N, S and H atoms are colored respectively in green, red, blue, yellow and gray. The toroidal patches and spherical polygons on fixed probes are colored respectively in orange and cyan. All the SES figures in this paper are prepared using our structural analysis and visualization program written in C++/Qt/OpenGL.

**Table 5:** The seven structures in 𝕊 with 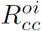 < 1.02. The nine SES-defined properties are the same as those in Table 3. The second column is the number of atoms in a structure. The last row lists their averages for 𝕊.

| pdbid | $n$ | $\rho_A$ | $\rho_B$ | $\rho$ | $\sigma_s$ | $\sigma_s^+$ | $\sigma_s^-$ | $R^{io}$ | $R_{cc}^{oi}$ | $A^{oi}$ |
| --- | --- | --- | --- | --- | --- | --- | --- | --- | --- | --- |
| 1he7 | 1615 | -0.02601 | 0.04437 | -0.00440 | -0.10921 | 0.16977 | -0.42917 | 1.61591 | 1.01392 | 1.52509 |
| 2ouw | 3989 | -0.02920 | 0.03028 | -0.00057 | -0.11053 | 0.17059 | -0.44484 | 1.46541 | 0.97992 | 1.73937 |
| 2qsk | 1269 | -0.02375 | 0.04558 | 0.00269 | -0.09007 | 0.17609 | -0.44169 | 1.50466 | 0.95139 | 1.31158 |
| 2rcd | 7809 | -0.03372 | 0.03441 | -0.00126 | -0.11634 | 0.17715 | -0.46242 | 1.51725 | 1.01543 | 1.46297 |
| 2x57 | 4655 | -0.03615 | 0.03399 | -0.00452 | -0.13306 | 0.17129 | -0.46970 | 1.68832 | 1.01039 | 1.32650 |
| 3qva | 6109 | -0.03236 | 0.02721 | 0.00034 | -0.11655 | 0.19231 | -0.45482 | 1.48751 | 0.99860 | 1.35483 |
| 3vqj | 3148 | -0.03425 | 0.03806 | -0.00226 | -0.11817 | 0.17355 | -0.46504 | 1.57286 | 0.98069 | 1.51827 |
| $\mathbb{S}$ | | **-0.0290** | **0.0270** | **-0.0015** | **-0.125** | **0.186** | **-0.490** | **1.875** | **1.496** | **1.214** |

A small *r*_*cc*_ for a single atom implies that it has a large *α*_s_ area, that is, the atom is much exposed to solvent and is thus a good candidate for hydrogen bonding if it is a polar atom. A small *r*_*cc*_ over a set of neighboring atoms means that the region formed by those atoms is locally-rugged and likely tightly-packed. Typically such a region has more accessible carbon atoms than a region with a larger *r*_*cc*_. In the contrary, a large *r*_*cc*_ for a single atom implies that the atom is largely hidden from the solvent while a large *r*_*cc*_ over a set of neighboring atoms means that they together form a locally-smooth surface region. Typically such a region has more accessible protons, oxygen and nitrogen atoms than a region with a smaller *r*_*cc*_. Compared with a rugged surface a smooth one is less disruptive to water’s hydrogen-bonded networks [7], and the VDW attraction between its surface atoms and solvent molecules is likely to be stronger. The proteins with a small *r*_*cc*_ have surface geometrical properties akin to those for the lipid-exposing regions of transmembrane proteins, lipid-protein, DNA-protein and PPI interfaces. VDW attraction has been shown to be important for the solvation of apolar molecules in aqueous solvent [17, 39]. As shown in Fig. 8 one salient feature of *r*_*cc*_ is that it increases with protein size but the rate of growth becomes smaller when the number of atoms in a structure is > 10,000. With more accessible atoms it becomes increasingly possible to form locally-smooth surface and consequently to have stronger VDW attraction between accessible atoms and solvent molecules. However it is obvious that some of the naturally-occurring proteins could remain soluble with a 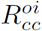 value close to 1.0 (Table 5) and there also seems to be an upper limit for 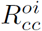 for all the naturally-occurring water-soluble proteins (Fig. 7 and Table 6). The limited 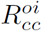 range for naturally-occurring water-soluble proteins suggests that their surfaces may have been optimized to interact with aqueous solvent. In the contrary some de novo designed proteins have rather large 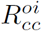 values (Fig. 7 and Table 6) [52] possibly because of the desire to enhance their solubility via so-called supercharging approach that increases the percentage of surface polar atoms over apolar ones. As shown in Fig. 10 the SES of a protein with a large 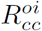 has largely-exposed *polar* atoms while a protein with a small 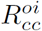 has largely-exposed *apolar* atoms.

**Figure 7:**
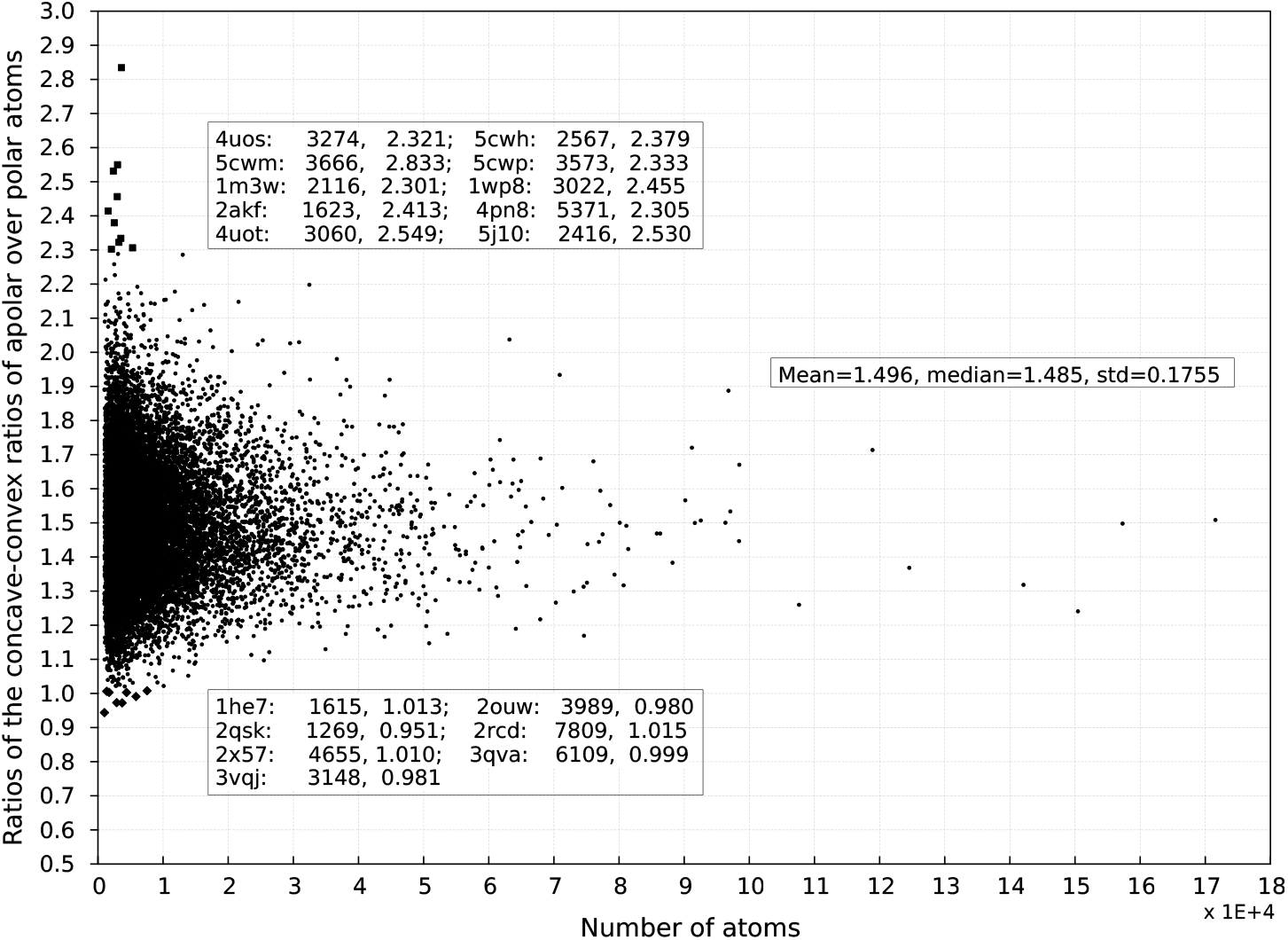
The 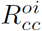s for 𝕊. The middle insert lists the mean, median and standard deviation for 𝕊. The top insert lists ten structures (depicted as filled squares) in 𝕊 that have 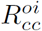 > 2.30. Among them are eight de novo designed proteins (4uos, 5cwh, 5cwm, 5cwp, 4pn8, 4uot and 5j10), a coiled coil protein (2akf) and a virus fusion core protein (1wp8). The bottom insert lists seven structures (depicted as filled diamonds) that have 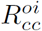 < 1.02. Among them are two PSI targets (2ouw and 2rcd) with unknown functions, a nerve growth factor binding site on Trka (1he7), a 5-hydroxyisourate hydrolase (3qva), a beta-carbonic anhydrase (3vqj) from thiobacillus thioparus, a polypeptide receptor (2×57) and an antiviral lectin scytoririn (2qsk) that has the smallest 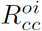(0.951) among all the proteins in 𝕊. The rest are depicted as filled circles. The x-axis is the number of atoms in a structure while the y-axis is 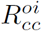.

**Table 6:** The ten structures in 𝕊 with 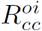 < 2.30. The nine SES-defined properties are the same as those in Table 3. The second column is the number of atoms in a structure. The first eight structures with the largest 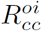s are de novo designed proteins. The last row lists their averages for 𝕊.

| pdbid | $n$ | $\rho_A$ | $\rho_B$ | $\rho$ | $\sigma_s$ | $\sigma_s^+$ | $\sigma_s^-$ | $R^{io}$ | $R_{cc}^{oi}$ | $A^{oi}$ |
| --- | --- | --- | --- | --- | --- | --- | --- | --- | --- | --- |
| 4uos | 3274 | -0.02290 | 0.05165 | 0.00392 | -0.07577 | 0.16608 | -0.47317 | 1.92180 | 2.32133 | 1.54527 |
| 5cwh | 2567 | -0.03282 | 0.03051 | -0.00674 | -0.15514 | 0.17671 | -0.54551 | 2.36745 | 2.37866 | 1.12571 |
| 5cwm | 3666 | -0.03443 | 0.03459 | -0.00309 | -0.15296 | 0.21979 | -0.58914 | 2.31827 | 2.83331 | 0.94308 |
| 5cwp | 3573 | -0.04159 | 0.04403 | -0.00620 | -0.16895 | 0.21358 | -0.58307 | 2.23531 | 2.33313 | 0.98420 |
| 1m3w | 2116 | -0.02827 | 0.04963 | -0.00059 | -0.10246 | 0.17904 | -0.48939 | 2.04249 | 2.30118 | 1.40759 |
| 4pn8 | 5371 | -0.01857 | 0.02348 | 0.00203 | -0.07430 | 0.18943 | -0.45380 | 2.16629 | 2.30531 | 1.15582 |
| 4uot | 3060 | -0.01570 | 0.02380 | 0.00332 | -0.08295 | 0.17818 | -0.47614 | 2.26879 | 2.54884 | 1.26948 |
| 5j10 | 2416 | -0.03282 | 0.03500 | -0.00357 | -0.13312 | 0.18631 | -0.54025 | 2.20735 | 2.53041 | 1.25704 |
| 1wp8 | 3022 | -0.03380 | 0.02995 | -0.00161 | -0.14487 | 0.18494 | -0.51638 | 2.24732 | 2.45540 | 1.06109 |
| 2akf | 1623 | -0.03772 | 0.05943 | -0.00396 | -0.15385 | 0.20559 | -0.56464 | 2.20482 | 2.41340 | 0.97170 |
| $\mathbb{S}$ | | **-0.0290** | **0.0270** | **-0.0015** | **-0.125** | **0.186** | **-0.490** | **1.875** | **1.496** | **1.214** |

**Figure 8:**
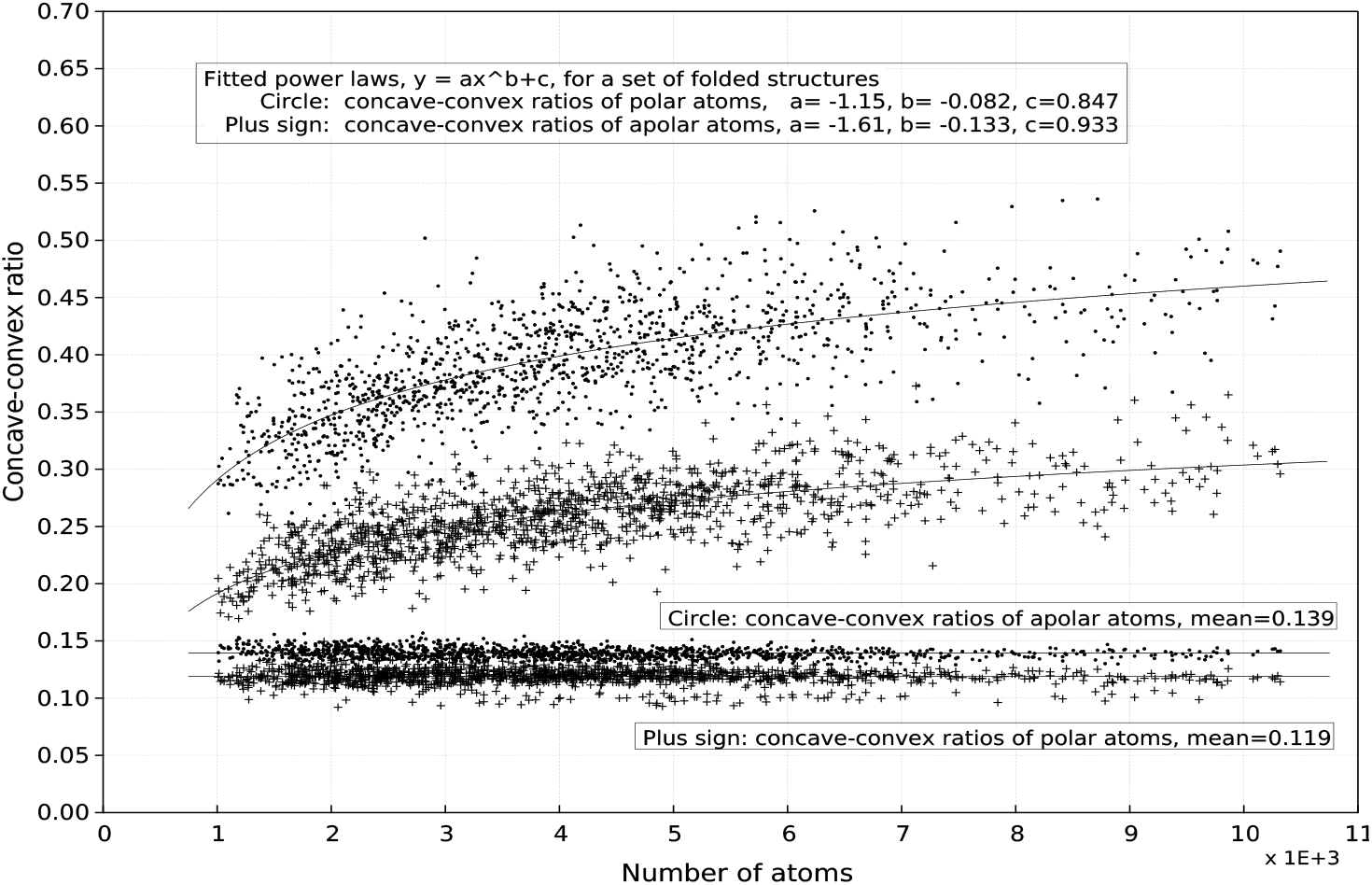
The concave-convex ratios (*r*_*cc*_s) over 𝕄_f_ and 𝕄_e_. The two curves represent respectively the fitted power laws for the 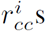 of the individual sets of the accessible polar atoms (plus signs) in 𝕄_f_ and for the 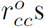 of the individual sets of the accessible apolar atoms (filled circles) in 𝕄_f_. The two lines indicate the respective means for the 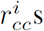 of the individual sets of the accessible polar atoms (plus signs) in 𝕄_e_ and for the 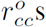 of the individual sets of the accessible apolar atoms (filled cirles) in 𝕄_e_. The x-axis is the number of atoms in a structure while the y-axis is *r*_*cc*_.

**Figure 9:**
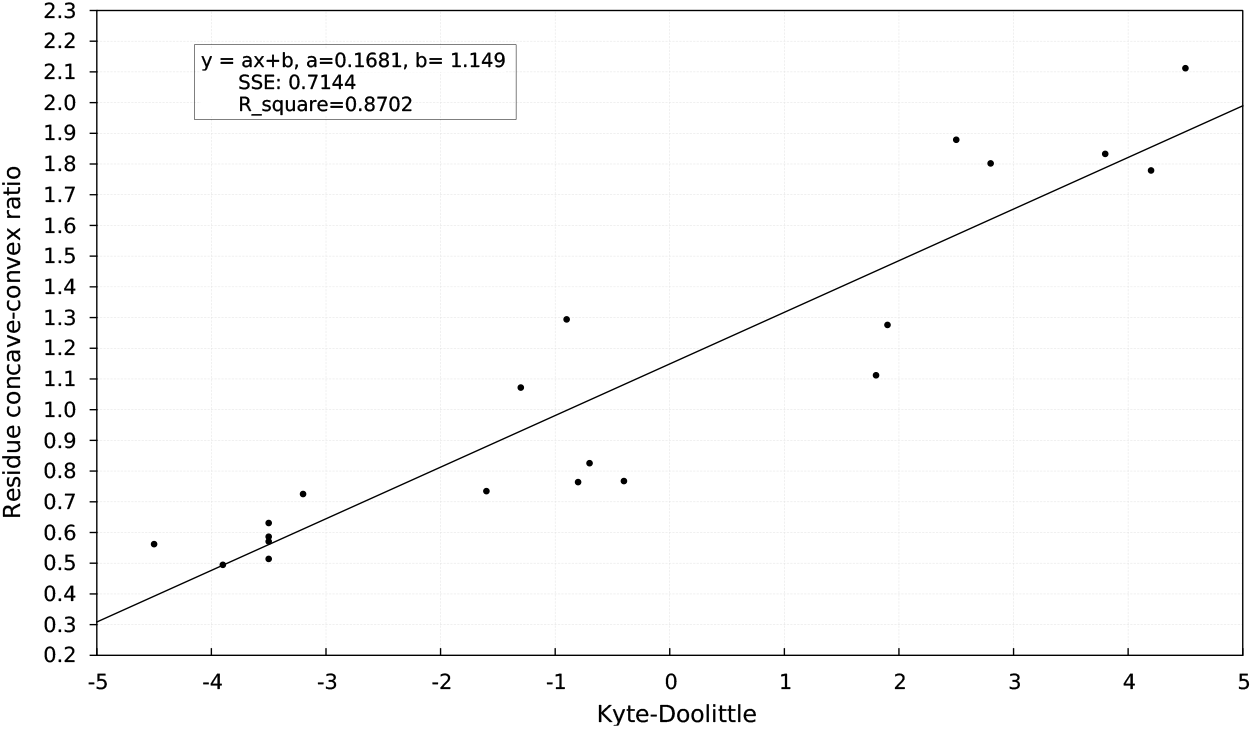
Twenty residue *r*_*cc*_s versus Kyte-Doolittle hydrophobicity scale. The line is a best-fitted linear equation. The x-axis is Kyte-Doolittle scale. The y-axis is residue *r*_*cc*_ average.

**Table 7:** Hydrophobicity scales versus accessible residue *r*_*cc*_ averages for 𝕄_f_. The five scales are respectively Kyte-Doolittle (KD) [42], Eisenberg-Weiss (EW) [43], Goldman-Engelman-Steitz (GES) [44], Janin (JANIN) [41] and experimental (EXP) hydrophobicity scales. The experimental data is from Table 2 of Kyte-Doolittle paper [42]. The correlations between accessible residue *r*_*cc*_ averages and the five scales are assessed by best-fitting them to a linear equation *y* = *ax* + *b* where *x* is hydrophobicity scale and *y* is residue *r*_*cc*_ average. The number in each cell is the coefficient of determination R_square_.

|  | KD | EW | GES | JANIN | EXP |
| --- | --- | --- | --- | --- | --- |
| Residue $r_{cc}$ | 0.87 | 0.71 | 0.51 | 0.90 | 0.63 |

**Figure 10:**
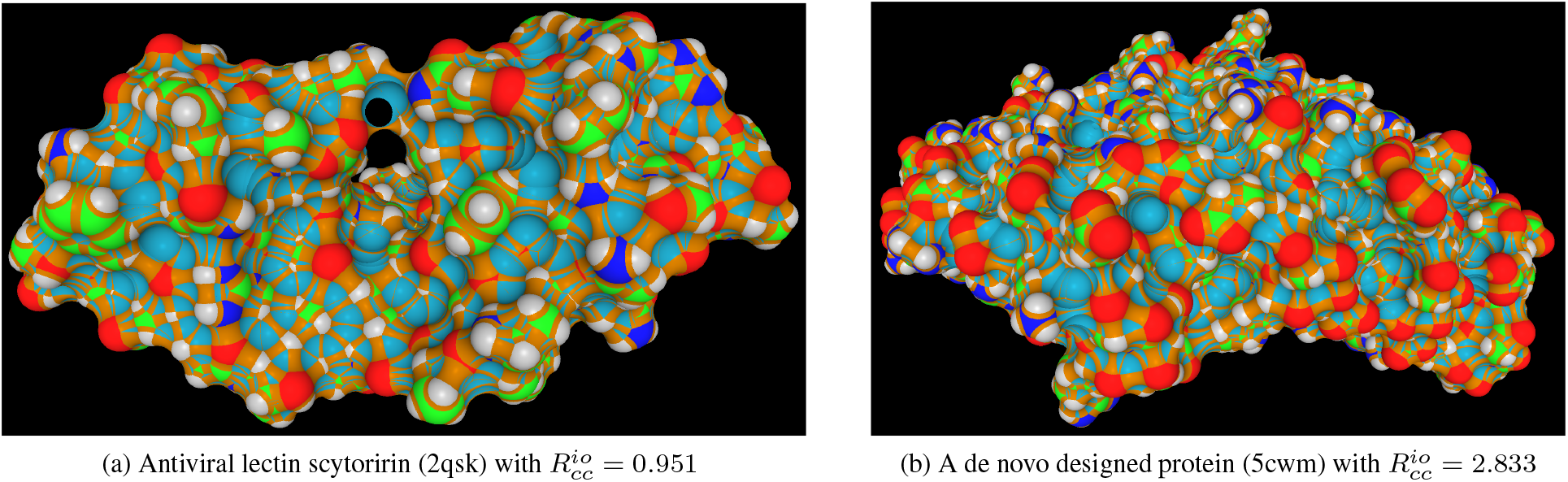
The SESs of the two structures with the smallest and the largest 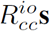 among all the structures in 𝕊. The coloring scheme is the same as in Fig. 6. The structure with the smallest 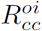 is a sugar binding protein with an antiviral activity while the structure with the largest 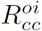 is a de novo designed protein [52].

In summary our large-scale analyses of the SES-defined properties *r*_*cc*_ and 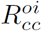 for different sets of protein structures and interfaces show that they likely pertain to protein solvation, folding and function possibly via the optimization of both the intermolecular VDW attractions between the accessible apolar atoms of a protein and solvent molecules and the intermolecular hydrogen bondings between its accessible polar atoms and solvent molecules. Since there exist no or only weak correlations between 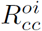 and SES-defined electrical properties *ρ* and *σ*_*s*_ (section S6 of the Supplementary Materials), 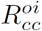 and likely *r*_*cc*_ are related more to intermolecular VDW attraction than to intermolecular hydrogen bonding. In addition the difference in *r*_*cc*_ between a surface apolar atom and a surface polar atom is in line with the heterogeneity of water motion in the first hydration shell. Taken together our SES analyses support the importance of VDW attraction to the solvation of an apolar molecule in a polar solvent [39].

### 3.5 Protein solubility and SES-defined properties

Previous analyses of the relationship between protein surface and protein-solvent interaction have focused mainly on SAS area and surface charge at residue-level [26, 31, 54, 34]. However, quantitative relationships between SESs and protein solvation and folding remain largely unknown and controversial [55, 56, 17, 57]. For example the past efforts to correlate SAS area with experimentally-determined solubility have only met limited success [9]. In the following we analyze two sets of experimental solubility data to illustrate the possible advantages of using atomic SES-defined properties to characterize protein-solvent interaction in general and protein solubility in particular.

#### 3.5.1 Experimentally-measured solubility and SES-defined properties

Recently Scholtz group has investigated seven proteins with crystal structures in order to find any correlations between experimentally-determined solubility and either SAS area or SAS-defined properties [9]. With the same goal we have analyzed the SESs of the same seven crystal structures with protons added by REDUCE [46]. As shown in Table 8 and Figs. 11 and 12, out of the list of SES-defined properties we have identified four of them that correlate well with the measured solubility data reported in their paper [9]. In the following we compare our SES-based analysis with their SAS-based analysis that uses only the SAS areas of heavy atoms since no protons have been added to any of the seven crystal structures. Though both our SES-based analysis (Figs. 11c, 11d, 12c and 12d) and their SAS-based analysis (Figs. 7 and 8 of their paper) have found good correlations between solubility and surface charge, important differences exist between the found correlations. Their SAS-based analysis had found only one good correlation with a R_square_ = 0.82 between the solubility in ammonium sulfate and the *absolute* value of net charge (Fig. 6F of their paper). In contrast, our SES-based analysis has found a good correlation with a R_square_ = 0.86 between Σ_A_ and solubility in ammonium sulfate (Fig. 11d) and a weak correlation with a R_square_ = 0.38 between Σ_A_ and solubility in PEG-8000 (Fig. 12d). In addition good correlations with respective R_square_ = 0.70 and R_square_ = 0.73 exist between 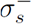 and solubility in both ammonium sulface (Fig. 11c) and PEG-8000 (Fig. 12c). In terms of SAS area, their SAS-based analysis had found good correlations with respective R_square_ = 0.81 and R_square_ = 0.84 between *fraction negatively-charged SAS area* and solubility in both ammonium sulfate (Fig. 8E of their paper) and PEG-8000 (Fig. 8F of their paper). As with their analysis we have found strong correlations with respective R_square_ = 0.84 and R_square_ = 0.94 between *R*^*io*^ and the solubility in both ammonium sulfate (Fig. 11a) and PEG-8000 (Fig. 12a). Most interestingly good correlations with respective R_square_ = 0.67 and R_square_ = 0.82 exist between *A*^oi^ and the solubility in both ammonium sulfate (Fig. 11b) and PEG-8000 (Fig. 12b). No similar correlations were reported in their paper [9]. The strong correlation between *R*^*io*^ and the solubility and the modest correlation between *A*^*oi*^ and the solubility sugget that the intermolecular hydrogen bonding interaction between accessible polar atoms and solvent molecules contributes largely to protein solubility. On the other hand, there exists no clear correlation between 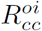 and solubility. Though the data set is rather small and thus the significance of these correlations is limited, the relevance to protein-solvent interaction of 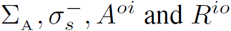 is consistent with the conclusions drawn from our large-scale SES analyses described earlier. And importantly these correlations between SES-defined property and protein solubility show that SES is better than or at least as good as SAS for the evaluation of surface area’s contribution to protein-solvent interaction in general and protein solubility in particular.

**Table 8:** The SES-defined properties for seven proteins with experimentally-determined solubility. The seven proteins are respectively *α*-chymotrysin (1yph), lysozyme (2vb1), human serum albumin (1e78), RNase Sa (1rgg), α-lactalbumin (1f6r), fibrinogen (3ghg) and ovalbumin (1ova). The six SES-defined properties are respectively net charge *Q,* the charge density of all the accessible atoms ∑_A_, the area-weighted charge density of the accessible atoms with negative partial charges 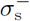, and three ratios *A*^*oi*^, *R*^*io*^ and 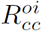. The last row lists their averages for 𝕊 except for *Q* where its range is listed. The second column is the number of atoms in a structure. The last two columns list respectively the experimentally-determined solubility values in ammonium sulfate (denoted as Log S_0_ * (NH_4_)_2_SO_4_ in their paper [9]) and PEG-8000 (denoted as Log S_0_ PEG in their paper) taken from Table 2 of their paper [9].

| pdbid | $n$ | $Q$ | $\Sigma_A$ | $\sigma_s^-$ | $A^{oi}$ | $R^{io}$ | $R_{cc}^{oi}$ | ammonium sulfate | PEG-8000 |
| --- | --- | --- | --- | --- | --- | --- | --- | --- | --- |
| 1yph | 6970 | 0.2559 | -0.00258 | -0.4797 | 1.0384 | 1.8987 | 1.3326 | 2.50 | 0.90 |
| 2vb1 | 1957 | 5.578 | -0.00113 | -0.4984 | 0.9286 | 1.8868 | 1.5773 | 3.50 | 1.50 |
| 1e78 | 16511 | -88.3123 | -0.00371 | -0.4342 | 1.6821 | 1.5956 | 1.3854 | 7.50 | 3.60 |
| 1rgg | 2881 | -14.3417 | -0.00290 | -0.4967 | 1.1502 | 1.8520 | 1.1706 | 3.40 | 1.60 |
| 1f6r | 11398 | -41.2857 | -0.00275 | -0.5016 | 1.1001 | 1.9009 | 1.5520 | 1.60 | 4.20 |
| 3ghg | 61292 | -51.4522 | -0.00277 | -0.4795 | 1.3104 | 1.7837 | 1.6303 | 3.50 | 1.70 |
| 1ova | 22101 | -38.7788 | -0.00313 | -0.4768 | 1.2926 | 1.7686 | 1.4006 | 6.00 | 2.00 |
| $\mathbb{S}$ | | **[-386.8, 127.5]** | **-0.002785** | **-0.490** | **1.214** | **1.875** | **1.496** | | |

**Figure 11:**
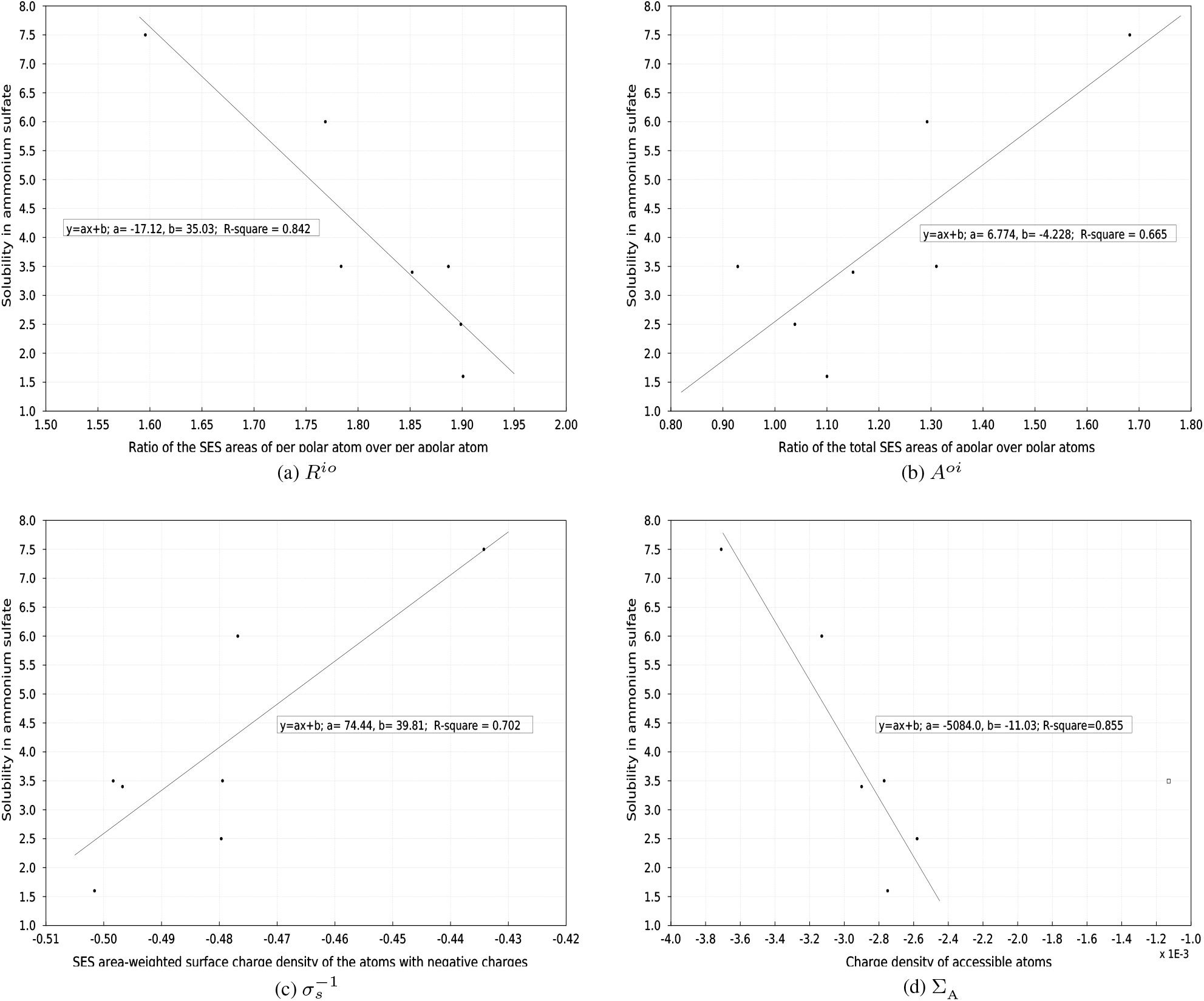
The four SES-defined properties 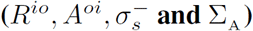 vs solubility in ammonium sulfate. The x-axes in figures (a, b, c, d) are respectively 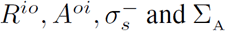 while the y-axis is the experimentally-determined solubility in ammonium sulfate. The inserted text in each figure shows the fitted linear equation and coefficient of determination R_square_. The data point for lysozyme (2vb1) depicted as an unfilled square is excluded in the fitting of Σ_A_ to solubility data.

**Figure 12:**
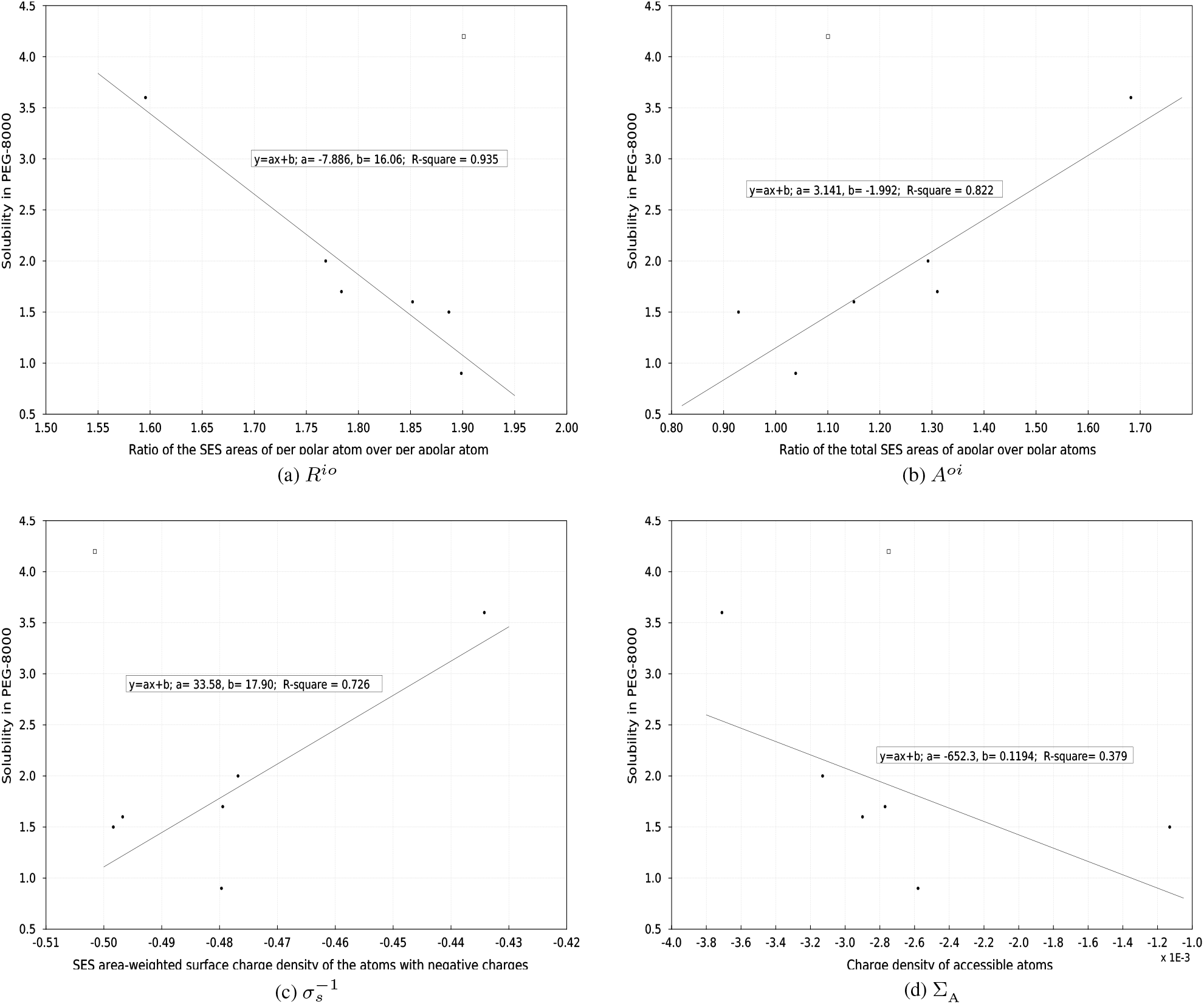
The four SES-defined properties 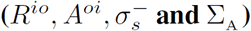 vs solubility in PEG-8000. The x-axes in figures (a, b, c, d) are respectively 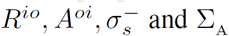 while the y-axis is the experimentally-determined solubility in PEG-8000. The inserted text in each figure shows the fitted linear equation and coefficient of determination R_square_. The data point for *α*-Lactalbumin (1f6r) depicted as an unfilled square is excluded in both our analysis and the SAS-based analysis [9].

#### 3.5.2 A water-soluble protein with a few titratable surface residues

In theory protein-solvent interaction is electrostatic in nature and thus the number of titratable surface residues in a protein is expected to be closely related to protein solubility. However, a recent protein redesign experiment by Winthers group [10] shows that the number of titratable surface residues in a protein is not a critical factor for its solubility. Specifically starting with a naturally-occurring protein (1exg) that has only four titratable surface residues (K28, D36, R68 and H90) Winther’s group has demonstrated that a soluble, functional protein with no titratable side chains could be engineered via protein redesign. It will be interesting to see whether the SES-defined properties for this particular protein differ largely from their averages for 𝕊. Since no structure is available for the redesigned protein and since the differences between 1exg and the redesigned one are likely to be small as far as their surfaces are concerned, we will compare the SES-defined properties for 1exg with those for 𝕊. As shown in Table 9 and Fig. S3 of the Supplementary Materials, except for the three *ρ*_*A*_, *ρ*_*B*_ and *ρ* that are somewhat more positive than their averages for 𝕊, the other six SES-defined properties, 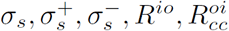 and *A^oi^,* are all rather close to their averages for 𝕊. In other words, at atomic level this particular protein is not an outstanding outlier in terms of the SES-defined properties that likely pertain to protein-solvent interaction. Thus 1exg and very likely the redesigned protein are expected to be as soluble as a typical protein in 𝕊 (section S5 of the Supplementary Materials). This example illustrates a possible advantage of SES-defined properties at atomic level over SAS-defined properties at residue-level for the description of protein solubility.

**Table 9:** The nine SES-defined physical and geometrical properties for 1exg. Only one negatively-charged residue (D36), two positively-charged residues (K28 and R68) and one histidine residue (H90) are solvent-accessible on the SES of 1exg. The nine properties are the same as that in Table 3. The last row lists their averages for 𝕊.

| | $\rho_A$ | $\rho_B$ | $\rho$ | $\sigma_s$ | $\sigma_s^+$ | $\sigma_s^-$ | $R^{io}$ | $R_{cc}^{oi}$ | $A^{oi}$ |
| --- | --- | --- | --- | --- | --- | --- | --- | --- | --- |
| 1exg | -0.016 | 0.027 | 0.0005 | -0.080 | 0.177 | -0.455 | 1.677 | 1.659 | 1.357 |
| $\mathbb{S}$ | **-0.029** | **0.027** | **-0.0015** | **-0.125** | **0.186** | **-0.490** | **1.875** | **1.496** | **1.214** |

### 3.6 The statistical distributions and power laws for SES-derived properties

At present the details of protein-solvent interaction could only be obtained through all-atom MD simulation with either explicit or implicit solvent models due to the amphipathic nature of the surface of a water-soluble protein. However long time all-atom MD with explicit solvent suffers from convergence problem especially for large-sized proteins while implicit models rely on *a prior* values for dielectric constants especially the dielectric constants near the surface of or inside a protein [16]. For example accurate dielectric constant for protein surface is the key for the computation of solvation free energy via electrostatic interaction. However the accurate determination of dielectric constants remains to be a challenging problem at present. As described above we have identified a list of SES-defined physical and geometrical properties that are likely to be important to protein-solvent interaction. Their statistical distributions and the power laws governing their changes with protein size obtained over large sets of high quality structures may help verify theories on anion solutes in protic solvent [12] or PLDL solvent model [58, 16]. In addition the statistical values and the power laws for SES-defined properties could be used to restraint the folding space of a protein and thus could serve as a term in an empirical scoring function for either protein structure prediction [59] or protein redesign [60, 61] or quality control in structure determination [62].

## 4 Conclusion

The solvent-accessible surface of a water-soluble protein is closely related to protein-solvent interaction and should have been adapted to the unique properties of aqueous solvent. To evaluate surface’s contributions to protein-solvent interaction and to find clues to surface’s adaptation to aqueous solvent we have analyzed the solvent-excluded surfaces (SESs) of four sets of water-soluble proteins and four sets of ligand-protein interaction interfaces. We discover that all the analyzed water-soluble proteins have a negative net surface charge. We have also identified a list of SES-defined physical and geometrical properties that are likely relevant to protein-solvent interaction based on their changes with protein size, their variations upon either unfolding or ligand-binding as well as the correlations between them and five known hydrophobicity scales and the correlations between them and experimentally-measured protein solubility. In contrast to previous structural analyses that focus mainly on accessible solvent surface area we find that surface charge is at least as important as surface area to protein-solvent interaction. Furthermore our analyses show that both the intermolecular hydrogen bondings between accessible polar atoms and solvent molecules and the intermolecular VDW attractions between accessible apolar atoms and solvent molecules contribute to protein-solvent interaction. These findings are consistent with water being a protic solvent prefers anions over cations and show that from a protein-solvent interaction perspective to fold into a native state is to simultaneously optimize net surface charge, intermolecular hydrogen bonding and VDW attraction rather than to only minimize apolar surface area. Our results suggest that the optimization of protein-solvent interaction through natural selection is achieved via (1) universal enrichment of negative surface charges for stronger intermolecular electrostatic interaction, (2) increased SES area for a polar atom for stronger intermolecular hydrogen bonding, and (3) higher concave-convex ratio for an accessible apolar atom for either stronger intermolecular VDW attraction or less disruption to solvent’s internal structure.

## Supplementary Materials

### S1: The polar atoms and apolar atoms in a protein

The list of polar atoms in a protein are:

K-HZ1, K-HZ2, K-HZ3, R-HE, R-HH11, R-HH12, R-HH21, R-HH22, Hp-HE2, D-OD1, D-OD2, E-OE1, E-OE2 HN, N, O, HT1, HT2, HT3, OT2, OT1 K-NZ, R-NE, R-NH1, R-NH2,N-OD1,N-ND2, N-HD21, N-HD22, Q-OE1, Q-NE2, Q-HE21, Q-HE22, He-ND1, He-NE2, He-HE2, Hd-ND1, Hd-HD1, Hd-NE2, Hp-ND1, Hp-HD1, Hp-NE2, Hp-HE2, H-ND1, H-HD1,H-NE2, H-HE2, S-OG, S-HG1, T-OG1, T-HG1, C-SG, C-HG1,Y-OH, Y-HH, W-NE1, W-HE1, Ep-HE2 and Dp-HD2.

The name of each atom consists of two parts separated by a hyphen: the part before the hyphen is residue name while the part after the hyphen is atom name in Charmm nomenclature. Each *polar* atom is either a hydrogen bond donor or an acceptor. The *apolar* atoms include all the other atoms in a protein.

### S2: The list of monomeric proteins in set 𝕄_f_ and set 𝕄_e_

1a58, 1a62, 1a76, 1a8d, 1a8l, 1a8q, 1a8s, 1aa0, 1ad6, 1ah7, 1ako, 1anu, 1ass, 1at0, 1avc, 1axn, 1azo, 1b0a, 1b8p, 1bgf, 1bhe, 1bkb, 1bm8, 1bqb, 1brt, 1bs2, 1byr, 1bz4, 1c1k, 1c44, 1cbg, 1cby, 1ciy, 1czt, 1d0b, 1d2p, 1d6m, 1dab, 1dd5, 1dd9, 1dhn, 1div, 1dov, 1dq3, 1dun, 1dus, 1dvo, 1dxh, 1dzf, 1e5m, 1e5w, 1ear, 1edg, 1edq, 1ee6, 1eg3, 1eok, 1eov, 1ep0, 1es5, 1et9, 1ew4, 1ezj, 1ezw, 1f2v, 1f82, 1fc6, 1fd9, 1fi4, 1fny, 1fob, 1fye, 1g2b, 1g43, 1g5z, 1g6a, 1g8a, 1g8p, 1g8s, 1g9g, 1gak, 1gen, 1gis, 1gpp, 1gqe, 1gs9, 1gvp, 1h4v, 1h6t, 1h6u, 1h7c, 1hjp, 1hq0, 1ht6, 1hus, 1hyq, 1i39, 1i4w, 1i5p, 1i60, 1ia6, 1idk, 1im4, 1im5, 1io0, 1io1, 1ipa, 1iq0, 1is1, 1iu9, 1iuh, 1iuz, 1ixh, 1ixk, 1ixl, 1j24, 1j27, 1j55, 1j7g, 1j8m, 1j93, 1jbk, 1jcf, 1jdw, 1jfx, 1jhc, 1jhs, 1jjf, 1jl5, 1jmm, 1jmw, 1jos, 1jrl, 1jvw, 1jyh, 1jyk, 1k04, 1k3v, 1k4n, 1k7i, 1k7j, 1kgs, 1kr4, 1ks5, 1ks8, 1l2f, 1l2l, 1lc0, 1lcy, 1lfp, 1lj5, 1ljo, 1lpj, 1lrv, 1lrz, 1ls1, 1lv7, 1lw3, 1ly1, 1lzl, 1m1h, 1m4l, 1mg4, 1mi8, 1mix, 1mug, 1muw, 1mw7, 1mw9, 1n67, 1nc5, 1nfn, 1ng6, 1nhy, 1ni3, 1ni5, 1nm2, 1nog, 1nq6, 1nri, 1nrw, 1nsj, 1nth, 1nty, 1o0x, 1o13, 1o73, 1o8p, 1o9g, 1oi7, 1ow1, 1ox0, 1ox3, 1oys, 1oyw, 1oyz, 1p1l, 1p1m, 1p2f, 1p3c, 1p4p, 1p7n, 1p99, 1pbj, 1pcs, 1pea, 1pgv, 1phz, 1pjr, 1psw, 1pv5, 1pvv, 1pvx, 1pyf, 1q2y, 1q5n, 1qcs, 1qhv, 1qme, 1qoi, 1qqe, 1qto, 1qw2, 1qyi, 1qz1, 1r3f, 1r5b, 1r6x, 1r8n, 1rh1, 1rh9, 1ri5, 1rjb, 1rl0, 1roc, 1rtt, 1ru4, 1ruw, 1rv9, 1rwz, 1rz2, 1s29, 1s2m, 1s2w, 1s2x, 1s48, 1s7i, 1s8n, 1scz, 1sdo, 1sek, 1sfs, 1sj8, 1sq1, 1sqh, 1sqw, 1srv, 1suj, 1sum, 1sur, 1syy, 1t1e, 1t1g, 1t5i, 1t6a, 1t71, 1t8k, 1t95, 1tev, 1tg0, 1thf, 1thx, 1tif, 1tjn, 1tn4, 1to3, 1tqg, 1tua, 1txd, 1txj, 1tyj, 1tzv, 1u14, 1u5h, 1u94, 1ub9, 1uds, 1uek, 1ujc, 1uku, 1uly, 1ulz, 1uok, 1usm, 1ux5, 1v05, 1v33, 1v43, 1v6t, 1v70, 1vaj, 1vbl, 1ve0, 1vgj, 1vgp, 1vyk, 1w0n, 1w5d, 1w8i, 1wch, 1wde, 1wdp, 1whi, 1wj9, 1wn2, 1wna, 1wos, 1wp5, 1wr2, 1wru, 1ws6, 1wu3, 1wv3, 1wvn, 1wwi, 1wxq, 1wy0, 1wza, 1wzz, 1x19, 1x3l, 1x7f, 1x9g, 1xdw, 1xdz, 1xeo, 1xeu, 1xhd, 1xip, 1xkr, 1xr5, 1xt0, 1xti, 1xub, 1xwl, 1xwy, 1y0k, 1y7e, 1y88, 1y8a, 1ydl, 1ydx, 1ye8, 1yfq, 1yh2, 1yhf, 1yii, 1yis, 1yle, 1ym5, 1ynm, 1yrv, 1ysp, 1yt3, 1yu0, 1yvr, 1yw5, 1yz6, 1z0w, 1z3x, 1z6m, 1z6n, 1z7h, 1z9l, 1zbm, 1zbp, 1zbs, 1zce, 1zcj, 1zd8, 1ziv, 1zjc, 1zma, 1zmr, 1zu4, 1zva, 1zxx, 1zyl, 2a4e, 2a4v, 2a6y, 2a6z, 2a9o, 2ae0, 2aeu, 2ah5, 2ahe, 2amh, 2amy, 2ap1, 2atr, 2au3, 2au5, 2au7, 2axq, 2b06, 2b0a, 2b18, 2b61, 2b8i, 2bdt, 2bep, 2bfw, 2bjq, 2bk8, 2bv6, 2bw0, 2bz7, 2c07, 2c08, 2c4n, 2c4x, 2cau, 2cc1, 2cgq, 2chr, 2ckw, 2cl3, 2cu2, 2cwp, 2cwy, 2cxc, 2cxh, 2cya, 2cyg, 2cyy, 2d4p, 2d4x, 2d58, 2d59, 2d5b, 2d7u, 2dbo, 2dfa, 2dg6, 2dg7, 2dh2, 2dp9, 2dpw, 2dvz, 2dwk, 2dxa, 2dyi, 2e01, 2e2c, 2e3u, 2e6m, 2efl, 2ehs, 2ejc, 2ek8, 2eo4, 2erf, 2et1, 2et6, 2ewf, 2f4q, 2f6h, 2f82, 2f9f, 2fb6, 2fbi, 2fc3, 2fd5, 2ffm, 2fg1, 2fi9, 2fl4, 2fm9, 2foz, 2fph, 2fq4, 2fu2, 2fy6, 2fzl, 2g29, 2g3a, 2g5x, 2gau, 2geb, 2ggo, 2glt, 2gq1, 2gs5, 2gs8, 2gsj, 2gwd, 2gwm, 2h1v, 2h36, 2h3g, 2h4r, 2h85, 2hbj, 2hhg, 2hm7, 2hrz, 2hvm, 2hz7, 2i49, 2i4a, 2i53, 2i5u, 2i6d, 2i6j, 2i6x, 2i7x, 2i88, 2ibl, 2ici, 2ict, 2idc, 2ii0, 2iih, 2ilr, 2iqt, 2iqy, 2j6b, 2ja2, 2jfr, 2nn5, 2nr7, 2nrj, 2nsa, 2nwh, 2nx2, 2nxc, 2nyv, 2o0m, 2o6q, 2o8l, 2o8n, 2obb, 2oca, 2ocz, 2odl, 2oeb, 2of3, 2ofz, 2ojh, 2oo2, 2ooe, 2op6, 2opj, 2oqr, 2ose, 2oy7, 2oyc, 2ozt, 2p0l, 2p17, 2p25, 2p2e, 2p4h, 2p51, 2p5d, 2p5i, 2p7n, 2pag, 2pbp, 2pcn, 2pge, 2ph1, 2pim, 2pjz, 2plc, 2pln, 2pn6, 2pom, 2pp6, 2ppn, 2psb, 2pth, 2pvu, 2q07, 2q0z, 2q13, 2q18, 2q4u, 2q5x, 2qa0, 2qc3, 2qc5, 2qff, 2qgm, 2qht, 2qi2, 2qip, 2qjl, 2qk1, 2qk2, 2qm3, 2qn0, 2qnk, 2qqy, 2qr3, 2qru, 2qsv, 2qv3, 2qwt, 2qx2, 2qy9, 2qyb, 2qyt, 2qyw, 2qyz, 2qz6, 2r48, 2r4g, 2r6q, 2r7j, 2r9i, 2ra1, 2rae, 2rbk, 2reu, 2rik, 2rjn, 2uu8, 2v5i, 2v8i, 2v9v, 2vaj, 2van, 2veq, 2vg9, 2vim, 2vk9, 2vpt, 2vri, 2vu5, 2w1n, 2w5q, 2w8n, 2wbx, 2wfb, 2wl1, 2wm8, 2wnx, 2x3m, 2x4l, 2xbt, 2xc2, 2xhc, 2xj4, 2xqh, 2xsa, 2xt0, 2xws, 2y5q, 2y6x, 2y9f, 2yci, 2yhs, 2ylm, 2yn0, 2yn2, 2yv2, 2yvy, 2ywe, 2ywj, 2ywk, 2ywr, 2ywx, 2yx5, 2z00, 2z01, 2z0m, 2z2u, 2z4u, 2z51, 2z5l, 2z6o, 2z7b, 2z8x, 2zcx, 2zeq, 2zhj, 2zrr, 2zxr, 3a2z, 3a3j, 3a7l, 3aam, 3ado, 3af5, 3agk, 3aj7, 3alf, 3aq1, 3asa, 3auf, 3av3, 3ayr, 3b02, 3b40, 3b43, 3b79, 3b7h, 3ba1, 3bb7, 3bbl, 3bh0, 3bjo, 3bjv, 3bk5, 3bkh, 3bn6, 3bod, 3bon, 3but, 3bw6, 3bwz, 3bz5, 3bzn, 3c12, 3c5v, 3c65, 3c7x, 3c8m, 3cax, 3cdi, 3cfz, 3chj, 3chm, 3ckm, 3cm0, 3cmi, 3cnu, 3cpe, 3csg, 3css, 3ctk, 3cwi, 3czc, 3d3a, 3d3y, 3d6l, 3d8m, 3dcy, 3dd4, 3dd6, 3deo, 3df7, 3df8, 3dgt, 3dnu, 3ds8, 3dsm, 3dso, 3du1, 3dyt, 3dz1, 3e0h, 3e13, 3e9l, 3ed5, 3eie, 3ejg, 3elx, 3enj, 3ers, 3etu, 3etv, 3eur, 3ewb, 3exc, 3exv, 3f4k, 3f67, 3fan, 3fbl, 3fbq, 3feu, 3ff2, 3fhf, 3fi7, 3fk8, 3foj, 3fp3, 3fqg, 3frr, 3ftd, 3ftj, 3fuq, 3fwt, 3fwu, 3fyn, 3fz4, 3g06, 3g40, 3g6s, 3g9g, 3ga2, 3gd6, 3gde, 3gha, 3go2, 3gr5, 3grh, 3grl, 3gs3, 3gt0, 3gx8, 3h04, 3h0x, 3h1g, 3h2g, 3h38, 3h6j, 3h6q, 3h7i, 3h7m, 3hcz, 3hdc, 3hdp, 3hjh, 3hp7, 3hpd, 3hr8, 3hra, 3hut, 3hvm, 3hvv, 3hvw, 3hxl, 3hz7, 3i32, 3i47, 3i8b, 3i9y, 3ibp, 3ic4, 3idv, 3ilc, 3ilv, 3im1, 3im8, 3im9, 3io0, 3ipc, 3ipz, 3iv3, 3iv4, 3ivf, 3jsr, 3jte, 3jv1, 3jxv, 3jyz, 3k01, 3k29, 3k5w, 3k63, 3k6i, 3k6j, 3k8w, 3kcw, 3kjh, 3kr9, 3kt9, 3ktn, 3kux, 3kwl, 3l3f, 3l4e, 3l7n, 3l8d, 3l9b, 3l9u, 3ld1, 3lda, 3lfp, 3lig, 3llb, 3lod, 3lop, 3lp5, 3lpz, 3lrv, 3lua, 3lx1, 3ly7, 3m16, 3m1e, 3m3h, 3m4x, 3m6c, 3m70, 3m7g, 3mah, 3mbr, 3mf6, 3mh7, 3mix, 3mm4, 3mpp, 3mtt, 3mx7, 3n26, 3n28, 3n2t, 3n3u, 3n4j, 3ne0, 3nf2, 3nft, 3nh4, 3nr5, 3ns4, 3o48, 3o59, 3o6p, 3o8z, 3oa7, 3obw, 3ohg, 3okq, 3oml, 3oop, 3or5, 3ozq, 3p51, 3p9n, 3pbi, 3pdd, 3pdg, 3pf9, 3pjx, 3pmm, 3pp8, 3pr9, 3ps5, 3psa, 3ptw, 3pwz, 3pyw, 3q3f, 3q69, 3q6l, 3q98, 3qav, 3qc7, 3qnm, 3qoo, 3qz6, 3r0r, 3r26, 3r2e, 3r2i, 3r2p, 3r38, 3r4c, 3r5e, 3r8q, 3rfy, 3rjp, 3rkg, 3rns, 3rrx, 3s4e, 3s8m, 3sbg, 3sft, 3sh4, 3shs, 3skq, 3slr, 3stp, 3sv0, 3sz7, 3t1w, 3t33, 3t5a, 3t8j, 3taw, 3tef, 3thi, 3tjt, 3tl2, 3tl4, 3tma, 3tpa, 3tqe, 3tql, 3tqo, 3tqq, 3tqz, 3trd, 3trg, 3ttg, 3txa, 3tyj, 3tys, 3u0r, 3u4k, 3u62, 3u97, 3ue3, 3ufb, 3ups, 3us6, 3utl, 3utn, 3ux2, 3v3t, 3v75, 3va9, 3vc5, 3vdg, 3vj9, 3vmn, 3vn5, 3vor, 3vub, 3vue, 3w1e, 3wa1, 3wap, 3wbi, 3whj, 3woh, 3wp4, 3wp8, 3wp9, 3wpa, 3wwa, 3wy8, 3zco, 3zpj, 3zqx, 3zsu, 3zyt, 4abl, 4ae7, 4amq, 4ams, 4anr, 4aur, 4axz, 4b0r, 4b8j, 4b96, 4b97, 4b9c, 4b9p, 4b9x, 4bin, 4btf, 4bwr, 4c3z, 4c7v, 4cbe, 4cfi, 4cg1, 4cil, 4cp6, 4cu2, 4cvr, 4cw4, 4dbd, 4dez, 4dh4, 4dhd, 4dim, 4dmv, 4dpb, 4e16, 4e22, 4e2u, 4e6h, 4e9l, 4eb0, 4ekz, 4es1, 4es6, 4esf, 4evf, 4ex6, 4ezb, 4f1r, 4f3q, 4f55, 4fbr, 4fcu, 4fd5, 4fmv, 4fnv, 4fs8, 4fwv, 4fx5, 4fzr, 4g0x, 4g2a, 4g2e, 4g3n, 4g54, 4g75, 4g9q, 4ga2, 4gb7, 4gbt, 4gco, 4gei, 4got, 4gou, 4gpr, 4grz, 4gzc, 4h60, 4h86, 4hbk, 4hcj, 4hd1, 4hde, 4hpn, 4hsp, 4htj, 4htl, 4hu2, 4hxt, 4i1t, 4i68, 4ic4, 4ic9, 4idh, 4idl, 4igi, 4ioy, 4iyk, 4izu, 4j4r, 4j4w, 4ja7, 4jcc, 4jg3, 4jmp, 4jp0, 4jwt, 4jz5, 4k1o, 4k2n, 4kds, 4kef, 4kg7, 4kpk, 4kqc, 4kqp, 4ksf, 4l0j, 4l0m, 4l1l, 4l4u, 4l6x, 4l8t, 4l9e, 4lcb, 4ldn, 4ler, 4leu, 4lf0, 4lgm, 4lj1, 4lkp, 4lmw, 4lru, 4lsw, 4ltt, 4lun, 4lzh, 4m9p, 4mag, 4me3, 4mfi, 4mh6, 4miw, 4mk6, 4mkx, 4mlw, 4mmh, 4mnr, 4mt7, 4mzd, 4n5a, 4n6q, 4nlm, 4nox, 4nux, 4o7i, 4o8b, 4ofx, 4ojm, 4oll, 4ovj, 4ox3, 4oy9, 4p09, 4p0l, 4p47, 4p48, 4p4u, 4p52, 4pau, 4pd0, 4ped, 4peu, 4pjr, 4pk9, 4pmh, 4pmx, 4ppu, 4ps6, 4pw0, 4pww, 4px8, 4py9, 4q62, 4q6b, 4q6v, 4q6x, 4q8r, 4qb7, 4qbo, 4qbu, 4qhe, 4qpr, 4qpt, 4qrl, 4qsg, 4quk, 4qvs, 4r0z, 4r5c, 4r5d, 4r6f, 4r6h, 4r6k, 4raa, 4rdb, 4rg8, 4rh4, 4rj9, 4rjz, 4rl1, 4rr5, 4rsf, 4rwu, 4u06, 4u4h, 4umi, 4uos, 4uu4, 4uvq, 4uw9, 4w5w, 4wcb, 4we2, 4wfi, 4wfv, 4wfx, 4wli, 4x1o, 4x2z, 4x9t, 4xh3, 4xof, 4xsj, 4xwx, 4xy3, 4y1w, 4y21, 4y23, 4y5j, 4y8f, 4yah, 4yj6, 4yn8, 4yno, 4yo1, 4z8z, 4z9x, 4zb3, 4zbh, 4zdm, 4zgi, 4zh0, 4zk3, 4zmi, 4zpj, 4zrx, 5a1m, 5a3y, 5aem, 5bp8, 5btb, 5bth, 5bxg, 5c86, 5coz, 5csm, 5dk6, 5e31, 5e43, 5efs.

### S3: The ratio of the number of apolar atoms over that of polar atoms

The ratio of the number of apolar atoms of a protein over its number of polar atoms *n*^*io*^ is a SES-defined property that could quantify the preference of polar atoms on its surface. As shown in Fig. S1 the *n*^*oi*^s for the proteins in 𝕊 increase very slowly with protein size and remain on average the same for large-sized proteins.

### S4: The ratio of the total area of apolar atoms over that of polar atoms

The ratio of the total area of apolar atoms of a protein over that of polar atoms *A*^*io*^ is a SES-defined property that may pertain to protein-solvent interaction. As shown in Fig. S2 the *A*^*oi*^s for the proteins in S do not change with protein size on average and the distribution with respect to the mean is symmetric with very close mean and median values.

### S5: The SES of a water-soluble protein with few titratable surface residues

In theory protein-solvent interaction is electrostatic in nature and thus the number of titratable surface residues in a protein is expected to be closely related to protein solubility. However, a recent protein redesign experiment by Winther’s group [10] shows that the number of titratable surface residues in a protein (1exg) is NOT a critical factor for its solubility. As described in the main text our analysis indicates that at atomic level in addition to surface charge and surface charge density other SES-defined properties may also contribute largely to protein solubility. As shown in Fig. S3 the SES for 1exg whose solubility has been studied by Winther’s group [10] is visually similar to a typical water-soluble protein in 𝕊.

**Figure S1:**
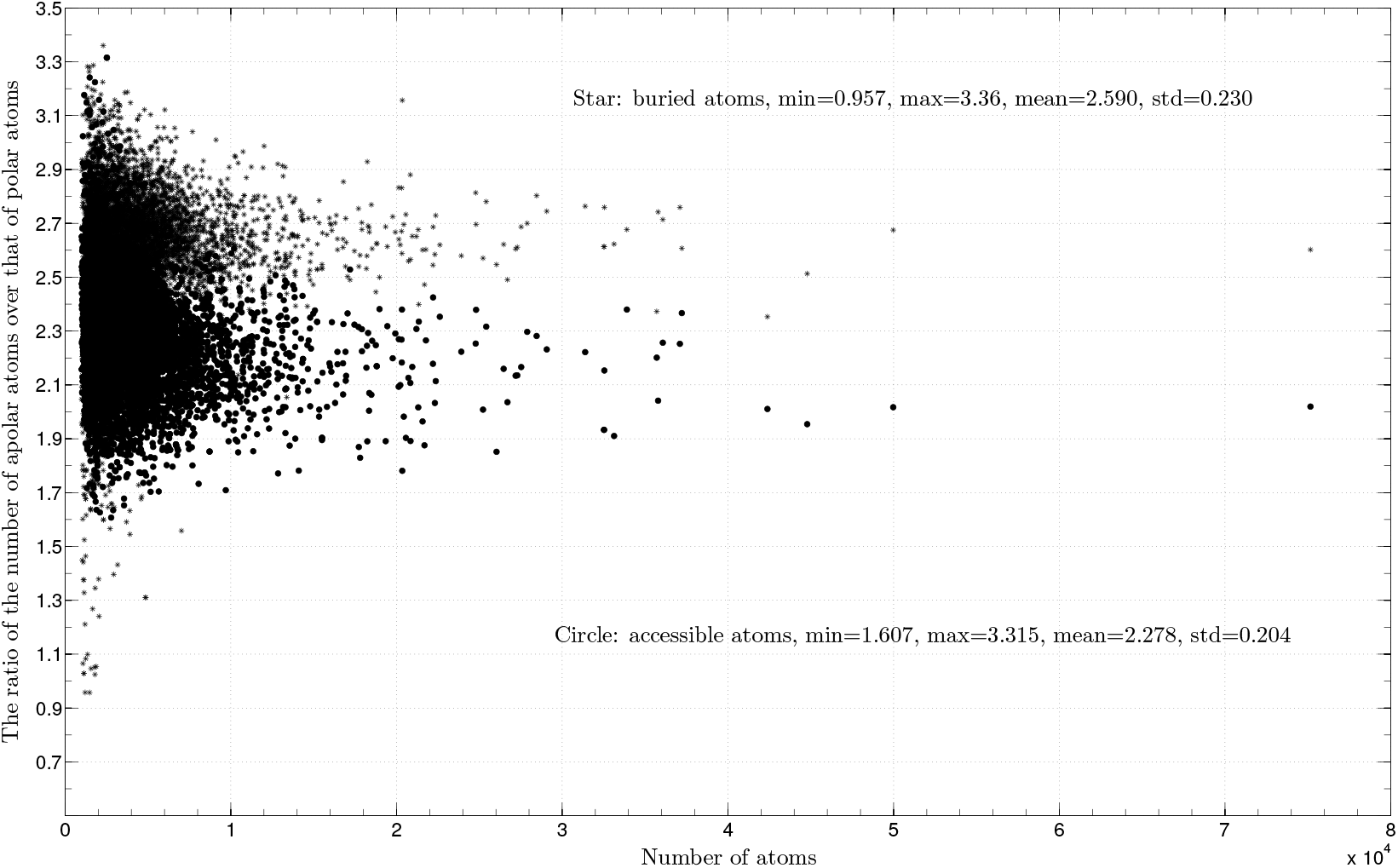
The *n^io^s* for 𝕊. The *n*^*io*^s for the individual sets of accessible atoms in 𝕊 are depicted as filled circles while the *n*^*io*^s for the individual sets of buried atoms as stars. The two inserts list their respective means and standard deviations. The x-axis is the number of atoms in a structure. The y-axis is *n^io^.* The 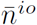 (2.590) for the buried atoms is 13.7% larger than the 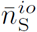 (2.278) for the accessible atoms.

**Figure S2:**
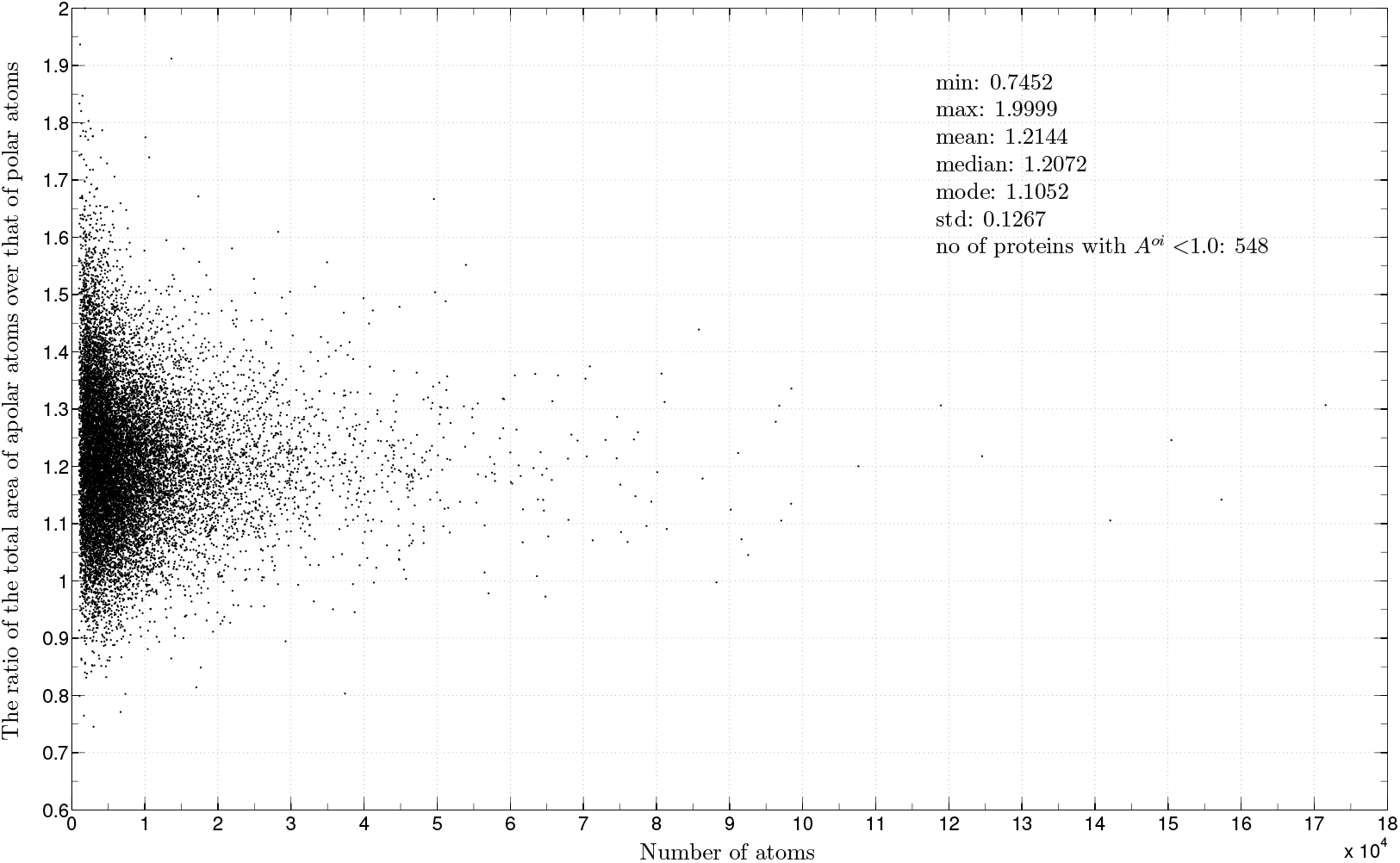
The *A^io^s* for 𝕊. The insert lists the minimum, maximum, mean, median, mode and standard deviation. There are 518 proteins in 𝕊 having *A*^*io*^ < 1.0. The x-axis is the number of atoms in a structure. The y-axis is *A^io^.*

**Figure S3:**
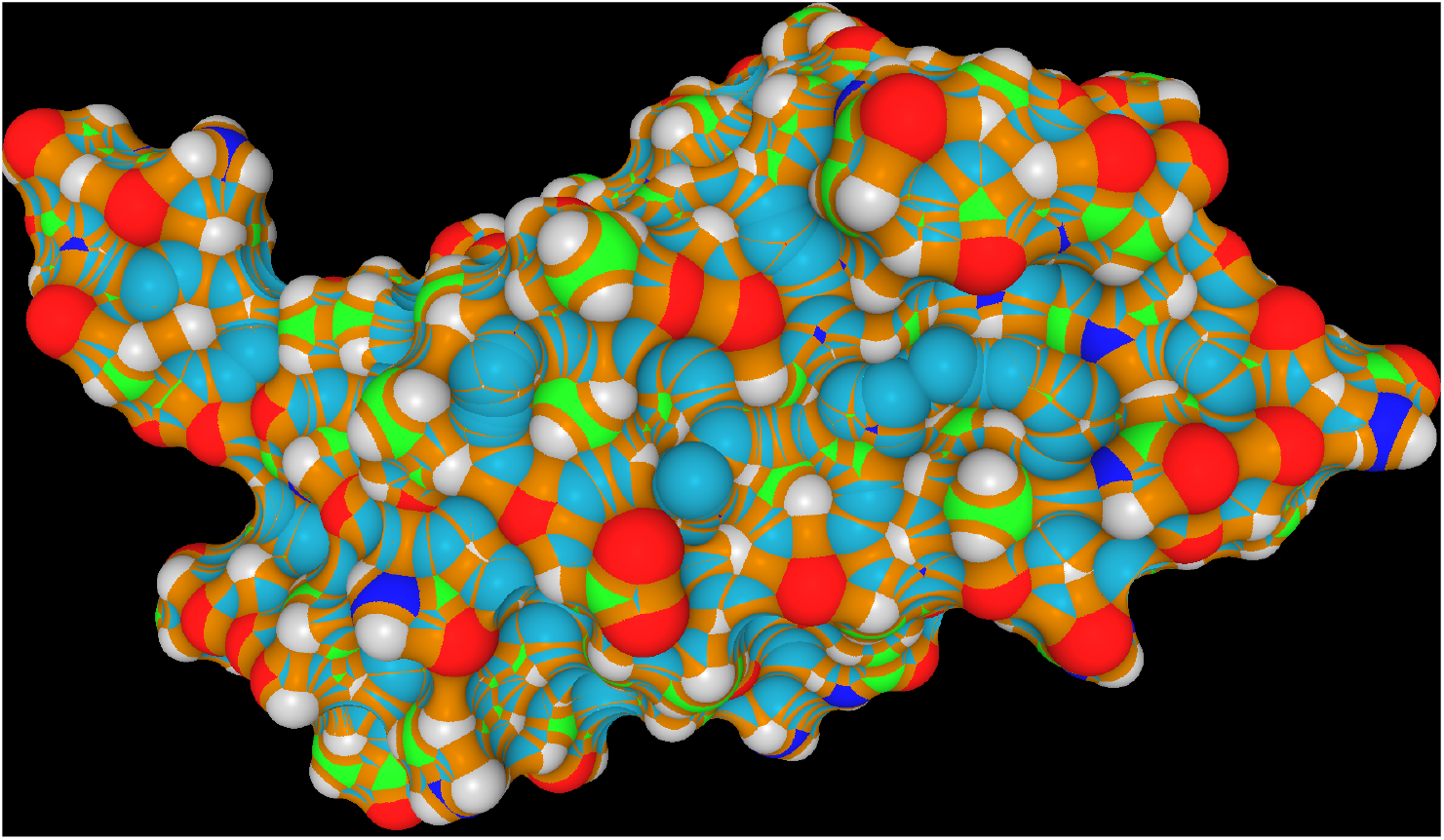
The SES of 1exg. There are only four titratable charged residues (K28, D36, R68 and H90) but many polar atoms on its SES. Consequently at atomic level its SES-defined physical and geometrical properties are close to their averages for all the water-soluble proteins in 𝕊. The coloring scheme for protein atoms is the same as Fig. 6 of the main text.

### S6: The independence between physical properties and geometrical properties of SESs

The preference of polar residues especially the charged ones on the surface of a water-soluble protein has been well-documented and frequently cited as a piece of evidence for the contribution to protein folding of hydrophobic effect. It has also been employed widely in de novo design of proteins to increase their solubility in aqueous solvent. As detailed in the main text the list of SES-defined physical and geometrical properties are closely related to either the electrostatic interaction or the hydrogen bonding interaction or the VDW interaction between surface atoms and solvent molecules and thus are relevant to protein-solvent interaction. Since all the three types of interactions are electrical in nature it is interesting to see whether the SES-defined physical properties are independent of the SES-defined geometrical properties. As shown in Figs. S4 and S5 there exist almost no correlations between the *R^io^s* and *ρ*_*A*_ s, between the *A*^*oi*^s and *ρ*_*A*_s and between the 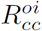s and *σ*_*s*_s for all the proteins in 𝕊. It implies that electrical property *ρ*_*A*_ is almost independent of geometric properties *A*^*oi*^ and *R*^*io*^, and electrical property *σ*_*s*_ does not correlate with geometric property 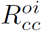. Since *ρ*_*A*_ and *σ*_*s*_ are defined in terms of surface charge while *A*^*oi*^, *R*^*io*^ and 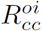 are the geometrical properties pertaining to hydrogen bonding and VDW interactions, the mutual independence of the former from the latter suggests that they evaluate protein-solvent interaction from different perspectives.

On the other hand, as shown in Fig. S6 there exists a modest correlation between the *R*^*io*^s and 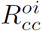s for the proteins in 𝕊 likely because both are defined in terms of **A**_i_ and **A**_0_ (Eqn. 7 of the main text).

**Figure S4:**
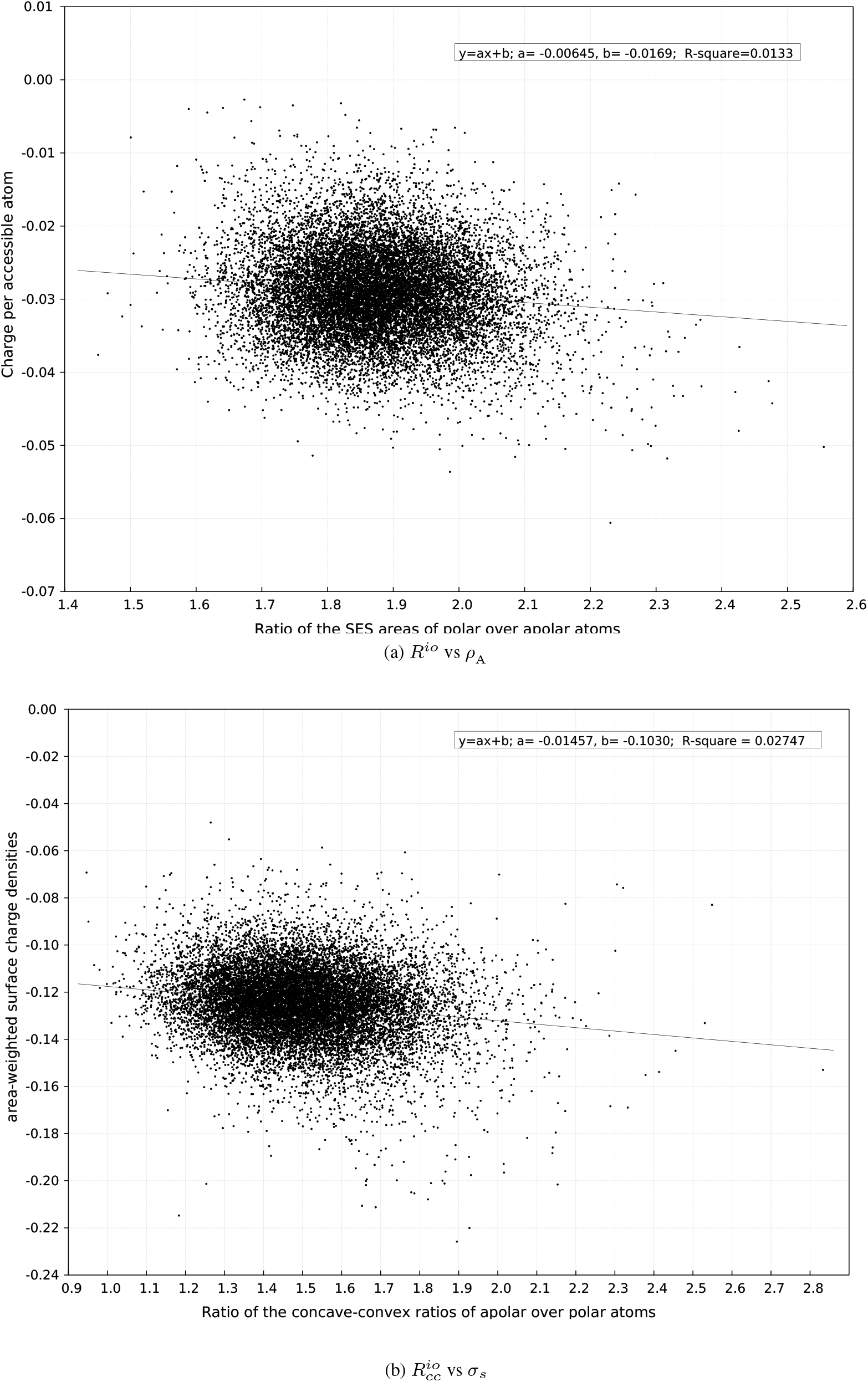
The independence of SES-defined geometrical and electrical properties. Figures (a) and (b) depict respectively the correlations between *R*^*io*^ and *ρ*_*A*_ and between 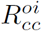s and *σ*_*s*_ for all the proteins in 𝕊. The two inserts list their respective fitted linear equations (shown as two lines) with coefficients of determination (R_square_s).

**Figure S5:**
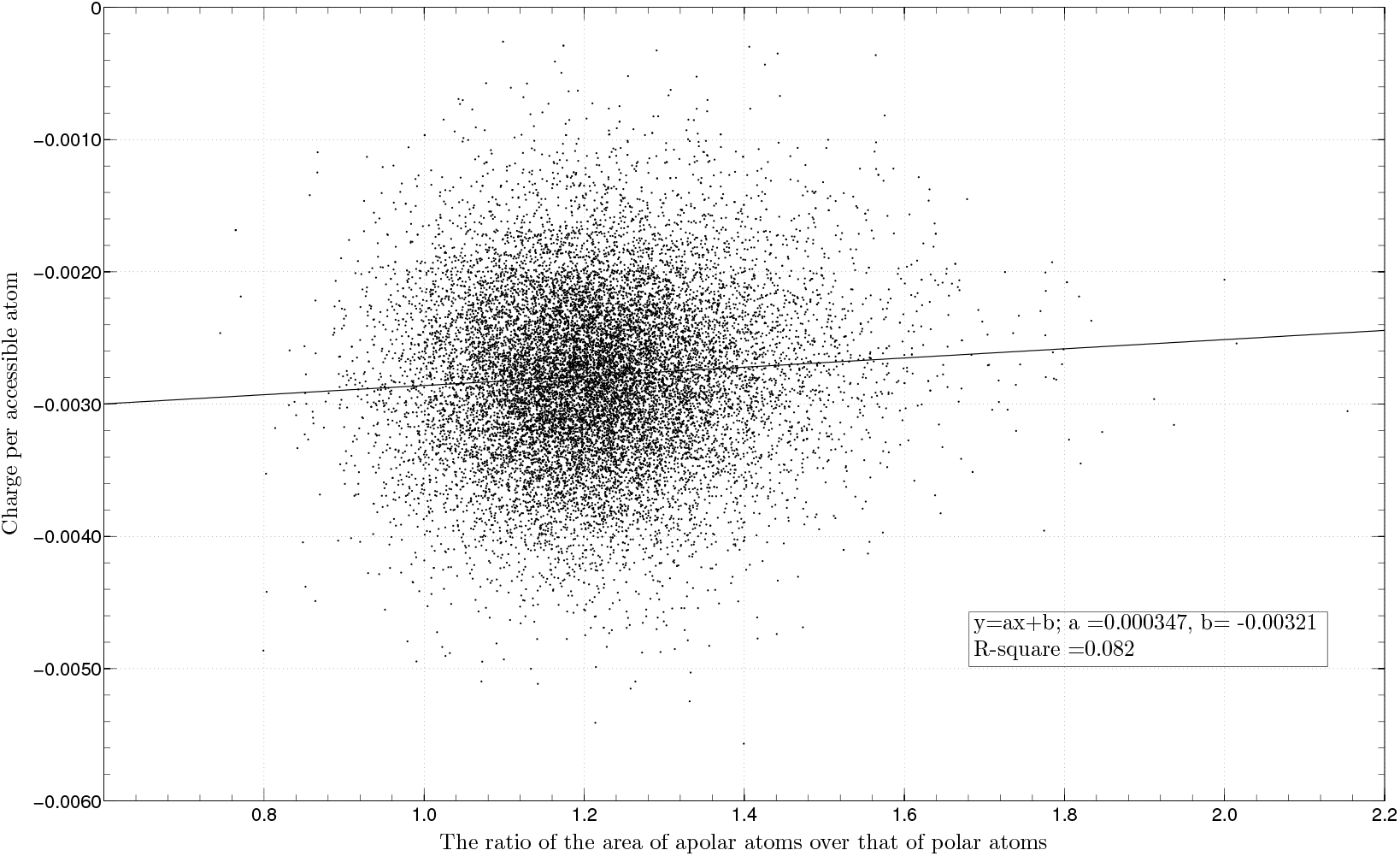
The independence of *A*^*oi*^ and *ρ*_*A*_. The inserted text lists the fitted linear equation with a coefficient of determination (R_square_ = 0.082). The x-axis is *A*^*io*^ while the y-axis is *ρ*_*A*_.

**Figure S6:**
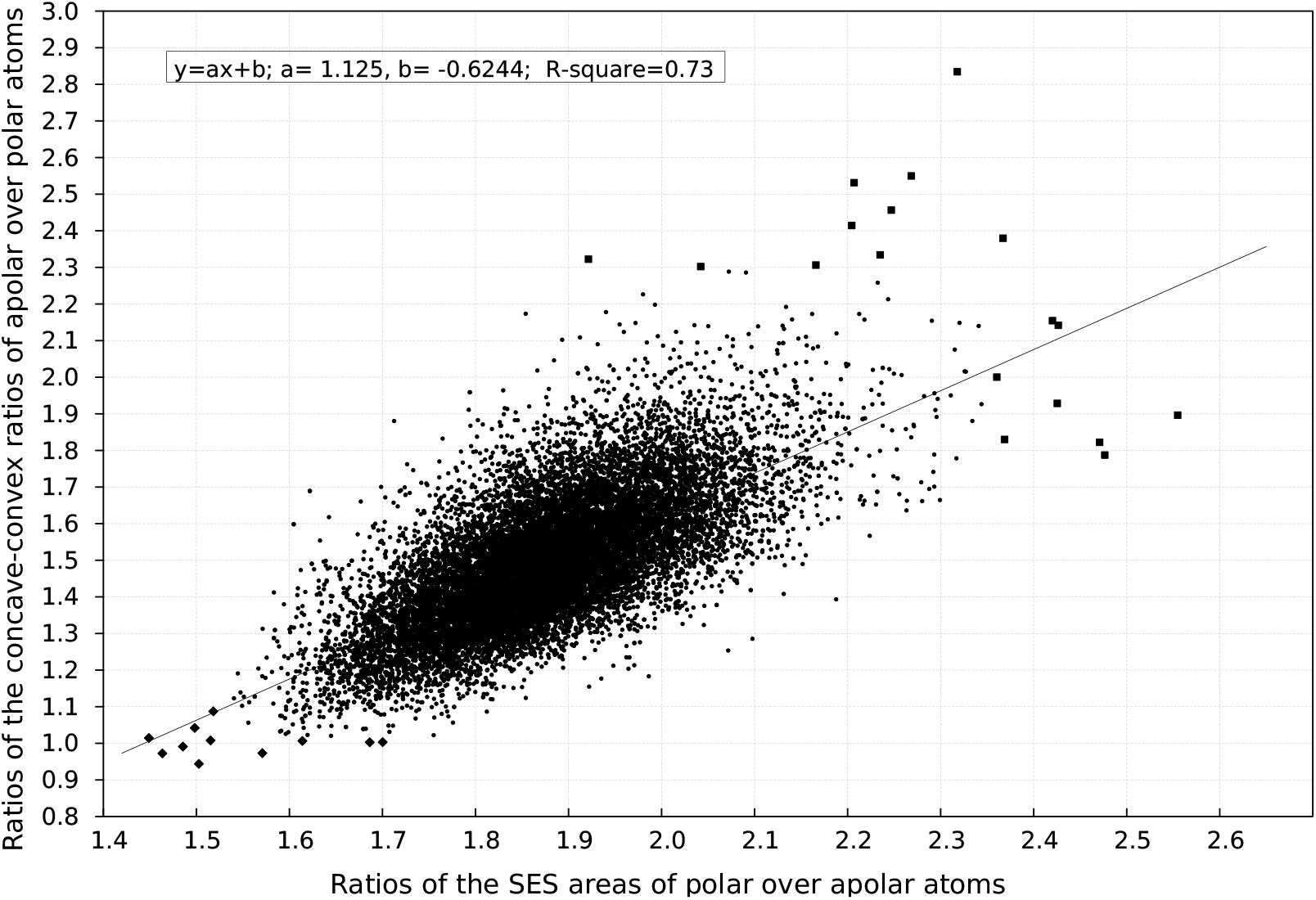
The correlation between *R*^*io*^ and 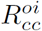. The inserted text lists the fitted linear equation with a coefficient of determination (R_square_ = 0.73). The x-axis is 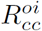 while the y-axis is *R^io^.* The structures in 𝕊 that have either *R*^*io*^ > 2.35 or 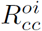 > 2.30 are depicted as filled squares while those that have either *R*^*io*^ < 1.54 or 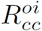 < 1.02 are depicted as filled diamonds. The rest are depicted as filled circles.

## Footnotes

1 Abbreviations: MD, molecular dynamics; SES, solvent-excluded surface; SAS, solvent-accessible surface; VDW, van der Waals; PPI, protein-protein interaction; DNA, deoxyribonucleic acid; 2D, two-dimensional; PDB, Protein Data Bank; PSI, protein structure initiative.

2 In the rest of the paper, *solvent-accessible atoms, accessible atoms, surface atoms* are used interchangeably.

3 In this paper intermolecular means between a solute and its solvent.

4 In this paper unfolding means the change from a folded structure to an extended conformation in 𝕄_e_ while the reverse change is called folding.

5 In terms of the list of SES-defined physical and geometrical properties described in this paper, no large differences exist between the SESs computed using 1.4Å probe radius and those computed using 1.2Å probe radius.

6 R. P. Feynman tried to explain the protein salt-out effect by assuming the existence of negative charges on protein surfaces.”The molecule (protein) has various charges on it, and it sometimes happens that there is a net charge, say negative, which is distributed along the chain”, The Feynman Lectures on Physics, page 7-10, Vol.2.

7 For a SES-defined property *x*, 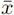 denotes its average over all the sets of accessible atoms in 𝕊 except for 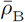 that denotes the average over all the sets of the buried atoms in 𝕊. For brevity such a 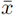 is to be written as either *x* average for the accessible atoms in 𝕊 or *x* average for the buried atoms in 𝕊 or simply as *x* average for 𝕊. The averages over 𝕄_e_ are to be written in the same manner.

8 In this paper protein size could mean either *n* or *n*_A_ or *A* since they are proportional to each other.

